# Generative design of sequence specific DNA binding proteins

**DOI:** 10.64898/2026.04.27.720408

**Authors:** Enisha Sehgal, Yuliya Politanska, Raktim Mitra, Paul T. Kim, Nayim González Rodríguez, Tushar Warrier, Andrew Kubaney, Akira Morishita, Riley Quijano, Jasper Butcher, Rohith Krishna, Robert J. Pecoraro, Brian Belmont, Nicole Roullier, Inna Goreshnik, Dionne K Vafeados, Paul Kwon, Rachana Ramarao, Jussi Taipale, Cameron J. Glasscock, David Baker

## Abstract

*De novo* protein design has advanced rapidly in recent years, yet the programmable recognition of specific DNA sequences remains a longstanding challenge. Here we describe a deep learning based approach for designing sequence selective DNA binding proteins. Our method combines structure generation using RFdiffusion3 with explicit screening against off-target interactions using AlphaFold3. We test this approach by generating 96 designs for each of 15 diverse DNA targets and identify specific binders for 7 targets, representing a ~100-fold improvement in success rates over previous approaches. We further characterize the binding landscape using variant competition assays and randomized library screening, revealing robust sequence discrimination across diverse targets. Together, these results represent a significant step forward in *de novo* sequence specific DNA binder design.

## Main

DNA binding proteins interpret and regulate the genome across all domains of life, enabling processes ranging from transcriptional control to genome maintenance ^1^. The ability to design proteins that recognize arbitrary DNA sequences would enable programmable gene regulation, targeted genome manipulation, and the creation of synthetic regulatory systems beyond those found in nature. Existing approaches to engineering DNA binding specificity largely rely on reprogramming native proteins such as zinc fingers ^2,3^ and transcription activator-like effectors ^4^, which, being family specific, constrain the accessible sequence space and structural diversity. The challenge of designing high-specificity protein–DNA interfaces *de novo* has largely limited deep learning efforts to predicting binding specificity rather than generating new binders ^5,6,7,8,9^, or language model-based redesign of native scaffolds ^10^. CRISPR-Cas9 takes advantage of nucleic acid base pairing to achieve DNA target selectivity, but its large size complicates delivery into cells. Additionally, the protospacer adjacent motif dependence restricts targeting space. Off-target binding and cleavage also remains a therapeutic challenge ^11^ with these systems. DNA binding proteins have also been engineered by computationally docking precomputed protein scaffolds onto target DNA structures and optimizing the resulting interface, but large library screening was required to identify sequence specific binders ^12^. A method for DNA binder design that can generate specific binders *de novo*, without requiring large scale screening is yet to be described.

We reasoned that diffusion-based generative methods, like RFdiffusion, could enable the custom generation of proteins optimal for binding specific target DNA sequences. These generative models have had considerable success in designing protein and small molecule binders, and proficient enzymes ^13,14^. While designed protein and small-molecule binders can achieve high target specificity, DNA presents a distinct challenge. B-form DNA adopts a similar global structure across sequences, and the chemical and geometric differences between bases are subtle, making sequence discrimination more difficult ^15,16,17^. We hypothesized that sequence specificity could be achieved efficiently by diffusion-based protein design, maximizing hydrogen bonding networks and packing interactions with DNA targets over extended regions (6-10 base pairs), while explicitly filtering out designs predicted to be non-specific.

## Design Method

We developed an RFdiffusion3 (RFD3) ^18^ based pipeline for protein-DNA binder design. A representative RFD3 diffusion trajectory around a DNA target is shown in Fig. 1a; starting from a Gaussian distribution offset from the center of the DNA target, diffusive denoising steps lead to spreading of the protein density to closely wrap around a 10 base pair region of the target structure. To generate a diversity of interaction modes, we sample a variety of placements of the protein center of mass relative to the DNA target using the RFD3 ori token feature, and a diverse set of hydrogen bond (Hbond) condition constraints (see Methods, Fig. 1b). Following RFD3 structure generation, we redesign the protein sequence using LigandMPNN ^19^, as we found that predictions of the structures of these sequences were closer to the designed complex structures than the original RFD3 generated sequences ^18^. Designs for which AlphaFold3 (AF3) ^20^ complex predictions were less than 8Å of the design model were subjected to an additional round of LigandMPNN sequence design and AF3 selection, this time imposing multiple cut-offs (AF3 confidence, structure alignment and protein-DNA interaction counts (see Methods). We found this sequence-structure prediction cycling, which roughly corresponds to one step of the optimization process used in the recently described ProteinHunter ^21^ and HalluDesign ^22^ protocols, substantially improved *in silico* metrics (Fig. 1b). We selected designs exhibiting dense atomic interactions with the target DNA to enable direct chemical readout of bases and phosphate backbone geometry^9^, as well as preorganized side-chain conformations at the interface, stabilized by intraprotein buttressing interactions to reduce off-target binding (see Methods). This binder generation pipeline is shown schematically in Fig. 1b and in more detail in Fig. S1.

**Figure 1.**
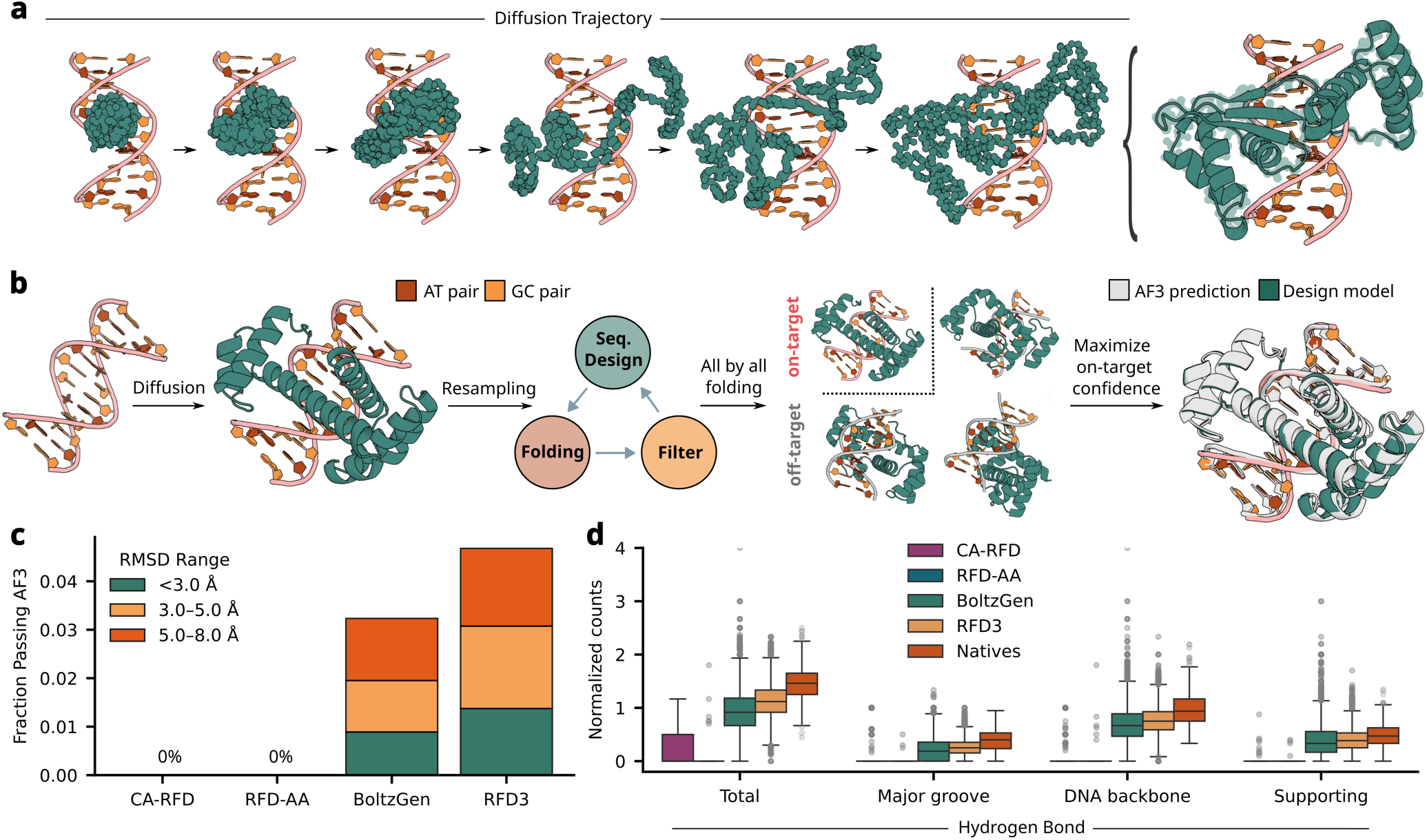
Design approach. **a**, Snapshots of RFD3 diffusion trajectory around a target DNA structure; only protein backbone atoms shown for clarity. **b**, Schematic of design pipeline. RFD3 is used to generate protein structures around a specific DNA target. The design is refined by multiple rounds of LigandMPNN sequence design, AF3 prediction and filtering, improving protein-DNA co-complex geometry (binder block, Fig. S1). This is followed by the specificity block (Fig. S1) which selects designs for which AF3 on-target predictions have higher confidence than AF3 off-target predictions. **c**, AF3 DNA-aligned protein RMSD pass rates compared across different generative models. DNA binder diffusion outputs with length randomly sampled from 120-150 amino acids (1000 backbones per target per model for core 6 targets (Table 1), 5 sequences per backbone with LigandMPNN). **d**, Interaction counts, per interacting nucleotide, across different generative models, calculated (using DSSR ^59^ version 1.7.8) on AF3 predictions post sequence design with LigandMPNN. The native protein set consists of 357 transcription factor PDB structures from the JASPAR database ^60^ with an average information content above 1.5.

To evaluate the performance of the RFD3-based pipeline *in silico*, we compared it to other generation methods: CA-RFdiffusion (CA-RFD) ^23^, RFdiffusion All-Atom (RFD-AA) ^24^, and Boltzgen ^25^. For each method, we carried out 1000 diffusion runs against six DNA targets (see Methods), and for each of the generated structures, we generated 5 sequences with LigandMPNN, followed by AF3 structure prediction. The atomic interaction based filtering and LigandMPNN-AF3 cycling steps were omitted for these comparisons. We found that the RFD3-based pipeline yielded designs with AF3 predictions more closely consistent with the design models (low DNA-aligned RMSD between design model and AF3 predicted structure) than the other methods (Fig. 1c). Boltzgen, which like AF3 is an all-atom diffusion model, was also considerably better than the residue based diffusion method CA-RFdiffusion, or the hybrid residue-atomic diffusion method RFdiffusion All-Atom. RFD3 and Boltzgen produced similar diversity of passing designs (Fig. S2). Detailed analysis of the designed protein-DNA interactions showed that the number of hydrogen bonds between protein sidechains and both the DNA bases and phosphate backbone was considerably higher using the all-atom diffusion methods compared to the residue based ones, and close to the density of atomic interactions in crystal structures of native transcription factors (TFs) bound to their targets (Fig. 1d, see Methods).

As noted above, a major challenge in designing DNA binding proteins is to achieve sequence specificity ^12^. While RFD3 generates designs interacting closely with the DNA target over an extended region, and making native-like levels of protein-DNA base hydrogen bond interactions, we cannot rule out the possibility that such interactions could also be made for other DNA targets, compromising sequence selectivity. We found that the minimum AF3-predicted aligned error (minPAE) among protein-DNA residue pairs recapitulated, to some extent, sequence selectivity of native transcription factor-DNA target complexes (Fig. S3); the minPAEs of AF3 predictions (with the protein templated and run in single-sequence mode) of TFs with their cognate DNA targets were in general lower than those with non-cognate targets (Fig. S3). We hence incorporated a final step into our design protocol in which designs were evaluated by comparing the AF3 confidence of predictions with the on-target DNA to those with off-target DNA sequences; designs were selected based on the difference between the on-target and best (minimum) off-target minPAEs (Fig. S4). In the remainder of the manuscript, we refer to the first phase of the design pipeline focused on binder design as the ‘binder block’, the second phase, as the ‘specificity block’ (Fig. S1), and the difference in the minPAE between target and off-target as the Δ*minPAE* (Fig. S4, see Methods).

## Experimental characterization

We used the above protocol to design binders to a core set of six DNA targets (Fig. S5) which contain no common 4-mers. 5 out of 6 of these targets are therapeutically relevant non-TF binding sites, and the sixth is the TATA box ^26^, for which no natural major groove recognizing binder exists. For each of these targets, at the end of the specificity block, we experimentally characterized the top 96 designs, ranked by Δ*minPAE*. In addition, to assess the impact of the specificity block and enable comparison with previous approaches lacking *in silico* specificity filtering, we experimentally characterized ~1,200 designs per target from the binder block alone. In the results presented below, designs from the specificity block are given the prefix ‘DBS’ and those from the binder block alone are named with the prefix ‘DBB’.

We used yeast surface display to evaluate binding of each selected design to each of the six target sequences. For the 96 specificity-block designs, genes were assembled individually from oligo pools, cloned into a yeast display vector, and transformed into yeast. Binding was quantified by flow cytometry as the normalized PE/FITC signal (DNA binding fluorescence relative to protein surface expression) in the presence of 1*µ*M fluorescent on-target or off-target DNA for each protein–DNA pair (Fig. 2a; Fig. S6, Methods). We considered a design specific if the normalized signal was higher for the on-target than for the five off-targets. Binder block designs were synthesized as an oligonucleotide pool and screened by fluorescence-activated cell sorting (FACS) followed by deep sequencing to identify designs enriched for their intended DNA targets with lower enrichment in off-target pools. For direct comparison with specificity block designs, individual clones were subsequently characterized using the same flow cytometry based assay.

**Figure 2.**
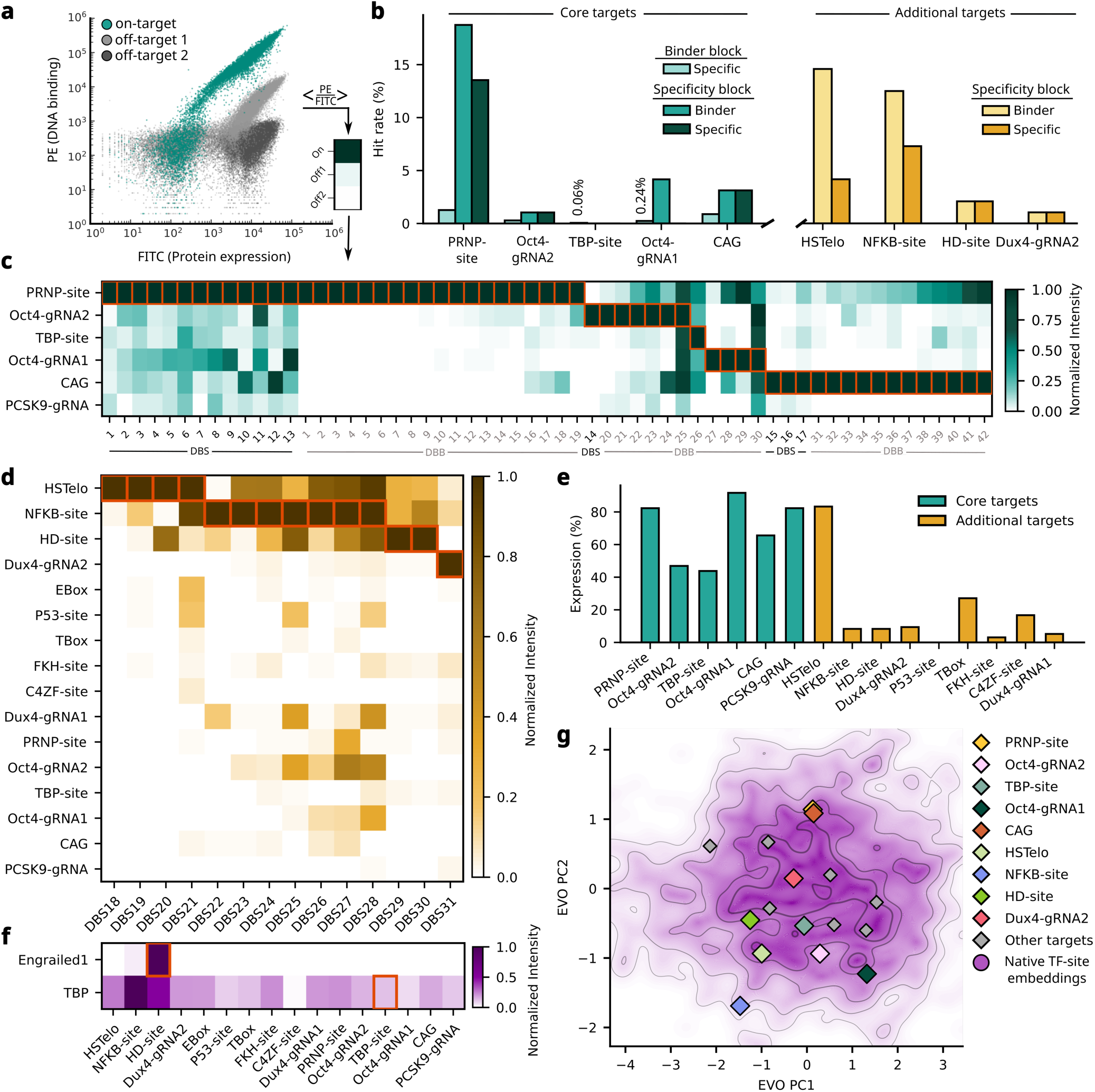
Experimental characterization of selectivity of designed DNA binding proteins. **a**, Overlay of representative flow cytometry traces for a single *de novo* DNA binder (DB), illustrating binding to on-target DNA (teal) and off-target DNA sequences (light and dark gray). Mean normalized PE/FITC values for each condition are used to generate the heatmaps shown in (c,d). **b**, Hit rates stratified by design category. Bars show the fraction of designs meeting binding and specificity criteria, grouped by binder block and specificity block libraries. For the specificity block, the binder hit rate (medium teal, light yellow) denotes designs with detectable on-target binding, whereas the specificity hit rate (light teal, dark teal, dark yellow) denotes designs with preferential binding to the on-target sequence (PE/FITC*_on_* > PE/FITC*_off_*). Targets without a successful specific design not shown. **c**, All-by-all specificity matrix for DBs against six core targets (Table 1), showing PE/FITC binding signal normalized by column. Designs were screened by yeast surface display at 1*µ*M DNA (non-avidity format). Red boxes denote the intended on-target sequence for each design. Gray labels indicate binder block designs; black labels indicate specificity block designs. **d**, All-by-all specificity matrix for DBs against additional targets (Table 1), displayed as in (**c**). **e**, Expression rates of specificity block designs measured by yeast surface display. Expression was quantified via FITC signal, with a threshold of <5-fold separation between low- and high-FITC populations used to classify expressing versus non-expressing designs. **f**, All-by-all specificity matrix for a subset of native transcription factors (TFs), displayed as in (**a**). Red boxes denote annotated target sequences derived from PWMs in the JASPAR database ^60^. Unipriot IDs for Engrailed1: P02836, and TBP: A0A7K4JSS6. **g**, Principal component analysis of Evo-derived embeddings of native TF binding sequences (see Methods), with *de novo* DB target sequences projected into the same space. PC1 and PC2 explain 63.2% and 19.8% of the variance, respectively.

The flow cytometry results for the most target specific designs obtained from the binder and specificity blocks are shown in Fig. 2c,d. Each column corresponds to an individual design (names at the bottom), and each row, to a different DNA target; colors indicate the normalized DNA binding signal and the red boxes, the intended target. A perfectly selective design, for example DBB1, has a strong binding signal for the intended target and little to no signal for other targets. From the binder block pools, we identified a total of 42 specific binders for 5 out of the 6 core targets, an average success rate of 0.46% over all binders tested. We found that the constraint *ΔminPAE* > 0 enriched for successful designs experimentally (Fig. S7). From the specificity block designs, we identified an on-target binder for 4 out of 6 core targets and a specific binder for 3 out of 6 core targets with success rates ranging from 1/96 (for Oct4-gRNA2) to 13/96 (for the PRNP-site) (Fig. 2b); the average success rate over all targets was ~3%. The *in silico* specificity filtering boosted the frequency of specific designs by 6 fold (comparing the binder block (0.5%) to the specificity block (3%)) and the combined protocol yielded specific designs at a rate 100 times higher than in our previous *de novo* DNA binder design efforts ^12^. Indeed, amongst the specificity block designs, most of the designs found to bind were also found to be specific (Fig. 2b). The specific designs exhibited high densities of protein-DNA inter-atomic interactions, consistent with native TFs and exceeding DNA binders generated by previous design pipelines (Fig. S8, Fig. S21). In total, we identified 59 specific binders (42 from binder block and 17 from specificity block) across 5 out of 6 core targets (Fig. 2c).

We next tested the generality of the design pipeline by making designs against ten additional targets spanning native TF binding sites, a human telomeric repeat (HSTelo) ^27^ sequence, and sequences adjacent to therapeutically relevant targets (Table 1). Unlike the six core target sites, these targets have some degree of sequence overlap, making the specificity challenge harder and more realistic (Fig. S5). Designs for each target were filtered *in silico* against all 15 off-targets, except for HSTelo which was only filtered against the six core targets. We obtained synthetic genes encoding 96 designs for 9 out of these 10 targets, selected in decreasing order of Δ*minPAE*, sampling up to 5 designs per diffusion backbone (the Ebox target was skipped due to low *in silico* pass rates, possibly due to symmetry effects). Flow cytometry characterization identified specific binders for 4 out of the 9 targets (HSTelo, the binding sites for homeodomain TF Engrailed, TF NF*ω*B, and the Dux4 gRNA site; Fig. 2d); roughly the same success rate found for the six core targets. Analogous to the specificity block results for the six core targets, most of the designs which bound their target were also specific: the binding rate ranged from ~1% to ~19% across the four targets while specificity rate ranged from ~1% to ~14%. (Fig. 2).

**Table 1.** Design targets. Summary of DNA target sequences evaluated in the de novo DNA-binding protein design campaign. Targets T1–T6 constitute the core set and were subjected to both the binder and specificity blocks. T7 was included in the specificity block but, during all-by-all in silico folding, was evaluated only against the core targets. Targets T8–T16 comprise additional sequences included in the specificity block and were evaluated against all targets in the all-by-all in silico analysis. TF target sequences were padded by random bases to reach length 10 or 12 (whichever is closer). MAXXXX.X entries in the source column denote JASPAR entries.

| Shorthand | Name | Sequence | Source | Family |
| --- | --- | --- | --- | --- |
| T1 | PRNP-site | TGAGGAGAGGAG | 61 | - |
| T2 | Oct4-gRNA2 | GGGCTTGCGA | - | - |
| T3 | TBP-site | CGTATAAACG | MA0108.3 | TATA-binding proteins |
| T4 | Oct4-gRNA1 | GGTGAAATGA | - | - |
| T5 | CAG | CAGCAGCAGCAG | 62 | - |
| T6 | PCSK9-gRNA | ACCTTCTAGG | 63 | - |
| T7 | HSTelo | AGGGTTAGGGTT | 27 | - |
| T8 | NFκB-site | GGGGATTCCCCC | MA0105.2 | RHR factors |
| T9 | HD-site | GCTTAATTAGCG | MA0094.2 | Homeo domain factors |
| T10 | Dux4-gRNA2 | CAGGCCGCAGG | 64 | - |
| T11 (no designs tested) | Ebox | ACCACGTGGT | MA0059.1 | bHLH |
| T12 | P53-site | AGACATGTCT | MA0106.3 | p53 domain factors |
| T13 | Tbox | AGGTGTGAAG | MA1919.2 | T-Box factors |
| T14 | FKH-site | GCGTAAACAA | MA0030.2 | Forkhead/winged helix factors |
| T15 | C4ZF-site | TGACCTTTGA | MA0017.1 | Nuclear receptors with C<sub>4</sub> zinc fingers |
| T16 | Dux4-gRNA1 | TCATCCAGCAG | 64 | - |

In total, through in-silico specificity filtering, we identified 31 specific binders across 7 different targets by testing only 96 designs per target, a significant methodological advancement in *de novo* DNA binder design. Combined with the binder block results, we identified specific binders for a total of 9 targets. These 9 targets span nearly as wide a range of sequence diversity as do the targets of native transcription factors (Fig. 2g), indicating the considerable potential of *de novo* design. The expression rate of designs filtered *in silico* against 15 off-targets were lower compared to designs filtered against 5 or 6 off-targets (Fig. 2e), suggesting a tradeoff between folding and selectivity ^28^ which will require further optimization in future design methods development. The amino acid sequences of the designs are completely unrelated to those of known proteins (Fig. S10), and the structures are globally very different from those of native DNA binding proteins (median TM-align against the PDB ^29,30^, of ~0.5, the cutoff for fold level structural similarity, see Fig. S11).

To compare the levels of specificity observed for our designs to those of native proteins under the same conditions, we sought to test 39 native DNA binding proteins recognizing sites in our target set in the same yeast display based assay. Only the engrailed homeodomain and the TATA bind protein (TBP) were well expressed on the yeast surface and bound their target (Fig. 2f, Fig. S9). The homeodomain was quite specific with slight off-target binding activity towards the NF*ω*B-site, analogous to our designed binder DBS29 targeting the same site (Fig. 2d). In contrast, TBP displayed considerable promiscuity (Fig. 2f), with high binding signal for several off-target sites; the short DNA targets used in our assay may be relatively malleable to the deformation this TF induces(PDB ID: 1TGH) ^31^. Thus, the specificity of our designs falls within the range observed for native DNA binding proteins under the conditions of the assay.

For more detailed experimental and affinity characterization, 82 of the designs were expressed in *Escherichia coli* and purified using immobilized metal ion affinity chromatography and size-exclusion chromatography (SEC). Of these, 46 were expressed solubly and were monomeric by SEC (Fig. S12). Biolayer interferometry (BLI) was used to determine K*_D_* for the cognate target site, which ranged from 3 to 30 nM (Fig. 3, Fig. S13). We also investigated the sequence specificity of a representative subset of the designs at single base resolution using yeast surface display based competition assay ^12^. For each single site variant, we measured the binding signal relative to the original target site. The results are shown in Fig. 4c, along with the overall structure of the design model (Fig. 4a) and a zoom in on the interactions within the highest specificity region (Fig. 4b); a schematic of the primary sidechain-base interactions is shown in ^32^ (Fig. S15). SELEX randomized library screening was also carried out for selected designs to assess specificity more globally (Fig. S14, see supplementary information). The following paragraphs summarize these data for a representative subset of the designs, relating the observed specificity to the designed structure and interactions.

**Figure 3.**
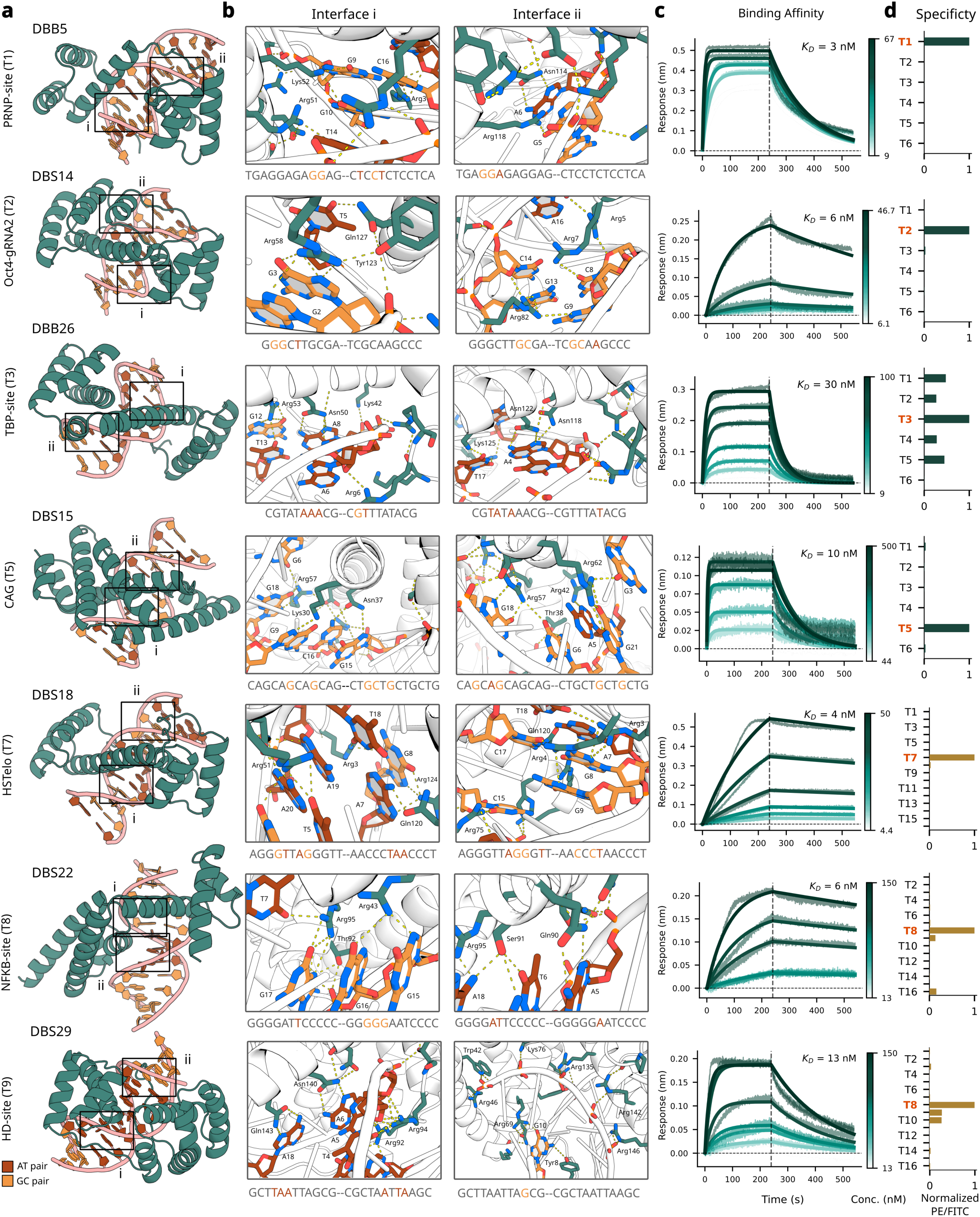
Structures and on-target binding affinities of selected specific designs. **a**, Designed protein–DNA complexes. Two interaction regions (interfaces i and ii) are indicated with boxes and shown in detail in (b). **b**, Close-up views of protein–DNA interfaces. DNA bases are colored by identity (A/T, red; G/C, orange), and key protein side chains are shown in teal. The full DNA target sequence is indicated below each panel, with interacting bases highlighted according to the same color scheme. Selected interactions are labeled. Base numbers in the reverse complement strand are offset by target length. **c**, Bio-layer interferometry (BLI) binding kinetics for selected designs. Sensorgrams show association and dissociation across a titration series of protein concentrations. Equilibrium dissociation constants (K*_D_*) were determined by global fitting to a 1:1 binding model using Octet Analysis Studio. **d**, Specificity profiles for selected designs, showing normalized binding signal (PE/FITC) across DNA targets. The intended on-target sequence for each design is indicated in orange on the y-axis (Table 1 for target annotations). Teal bars represent hits from the core target set and yellow represents the hits for the additional target set.

**Figure 4.**
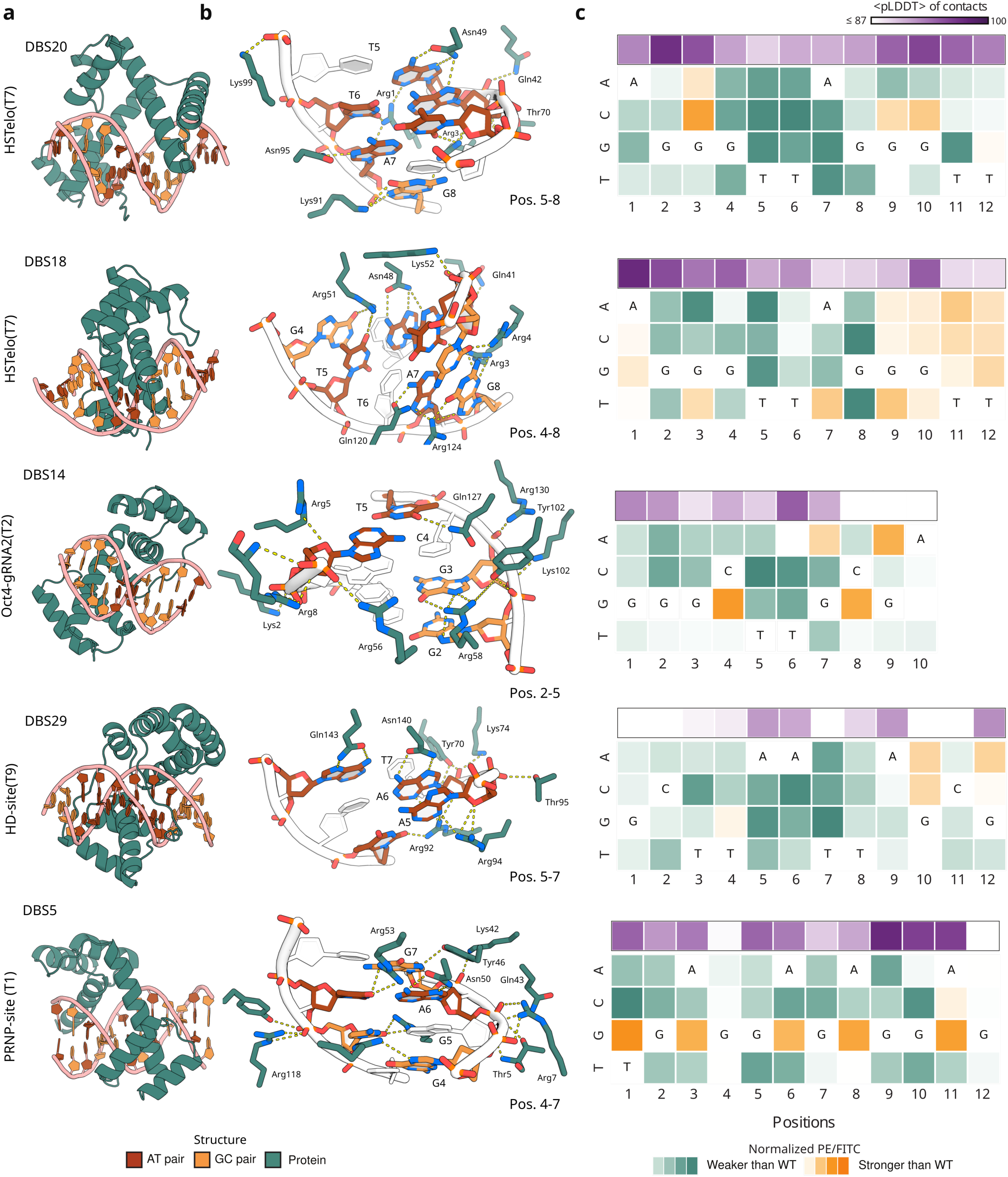
Characterization of specificity at single base resolution. **a**, Design models (AF3 predictions) of selected designs characterized for fine-grained specificity using yeast surface display-based competition assay, assessing binding towards all possible single nucleotide variants of corresponding DNA target. **b**, Protein-DNA contacts of selected regions shown per design. Nucleotides with no non-phosphate interaction are in white. Positions are annotated as per panel (**c**). **c**, Results of competition assay (normalized PE/FITC) for each design (orange-green heatmap) presented in context of average AF3 pLDDT of contacting protein residues (within 4Å) per base-pair position (purple heatmap). Green indicates a mutant competitor target was weaker than the intended (WT) base at that position, while orange indicates higher binding for the mutant competitor.

Design DBB5 (Fig. 3, row 1) against the almost fully poly-purine PRNP-site (Table 1) contains two helical subdomains closely interacting with different regions of the DNA target that are connected by an extended region not in regular secondary structure. Zoom-ins on the interacting regions (columns 2 and 3; labeled i and ii in the overall view in the first column) show detailed sidechain-base interactions including bidentate interaction of Asn114 specifically selecting A6. BLI affinity measurements (Fig. 3c) indicate binding affinities of 3 nm for this design. DBB3, which targets the same site, has a binding affinity of 10 nm (Fig. S13). HT-SELEX on DBB3 shows (Fig. S13) specificity towards the 5-mer GAGGA (repeated twice in the target) (Fig. S14). DBS5, a specificity block design (Fig. 4a, row 5) targeting the same site also reveals a clear preference for polypurines (through variant competition assay), although guanine is preferred over adenine (Fig. 4c row 5), likely due to an abundance of arginine sidechains present in the interface (Fig. S15b, row 5). Over positions 1-10, DBS5 prefers the intended target sequence over 35 out of the 40 single base variants tested in the competition experiments showing considerable specificity (Fig. 4c, row 5). Neither the AF3 minPAE (Fig. S17) nor geometric deep learning based specificity prediction methods ^8,9^ (Fig. S16) were able to predict the behaviour of this design under variant competition assay.

In DBS14 (Fig. 3, row 2) against the Oct4-gRNA2 site (Table 1), Arg58 in interface i of DBS14 makes a bidentate interaction with G3 (likely contributing to specificity ^33^, and is buttressed by a hydrogen bond from adjacent Tyr123. Tyr123 also makes a supporting polar contact with Gln127, which recognizes T5 via hydrogen bonding; such preorganized sidechain interactions are a product of our filtering strategy. Hydrogen bonds by positively charged Arg82, Arg7 and Arg5 to multiple bases in the minor groove (interface ii) likely contribute to both affinity and specificity ^34,35^. These interactions collectively make DBS14 quite specific against single nucleotide variants (Fig. 4, row 3). Affinity measurements for this design show low 6 nM binding affinity (Fig. 3c, row 2). DBS14 (Fig. 4a, row 3) is quite specific towards the target Oct4-gRNA2 at positions 1 to 7 with only 2 out of 28 mutations having stronger signal. This is driven by multiple base-specific major groove hydrogen bonds (T5 via Gln127, G3-G2 by Arg58 and Tyr102; Fig. 4b, row 3) and other interactions (Fig. S15, row 3).

Design DBB26 (Fig. 3, row 3) targets the AT-rich TATA box sequence (Table 1) via multiple base specific polar contacts in the major groove. A8 and A4 are recognized by bidentate amine sidechains Asn50 and Asn122 (interface i and ii), which are buttressed by other polar sidechains. A dinucleotide GT step (G12-T13) is also contacted by Arg53 (interface i). Whereas the native TATA-binding protein (PDB: 1TGH) which recognizes the same site primarily interacts with the DNA minor groove, DBB26 interacts primarily with the major groove, indicating that major groove recognition with high affinity (30 nM, Fig. 3c, row 3) and on-target selectivity (Fig. 3d, row 3) is possible for this sequence.

DBS15 (Fig. 3a, row 4) targets the CAG repeat sequence (Table 1) through extensive hydrogen bond networks and an integrated extended single domain topology. Asn37 in interface i hydrogen bonds with a dinucleotide G15-C16 step and is buttressed by a polar contact network involving other sidechains (interface i). In interface ii, Arg47 forms bidentate interactions with G18, and Arg62, with G21. Multiple hydrogen bonds to DNA phosphates likely contribute to affinity and stabilize the bound conformation of the DNA backbone, facilitating indirect readout of the target sequence ^36^. These interactions lead to an affinity of 10 nM (Fig. 3c, row 4) and strong on-target selectivity (Fig. 3, row 4). DBB32, another design towards the same target strongly recognizes the repeating unit GCA Fig. S14), overlapping with the reverse complement repeating unit TGC (which is a GCA offsetted by one base in the opposite strand, Fig. S14) indicating potential dimeric or multimeric binding.

DBS18 (Fig. 3, row 5) targets the human telomeric repeat sequence (HSTelo) (Table 1), contacting a G4-T5 step with Arg51 and A7-G8 step with Gln120 and Arg124 (interface i). This design also features a loopy extension into the minor groove (interface ii) housing multiple positively charged side chains in the negatively charged environment. Perhaps because of these electrostatic interactions, the design has a strong 4 nM affinity for the target (Fig. 3c, row 5) while still being considerably sequence selective Fig. 3d, row 5. Variant competition assays show (Fig. 4) the loopy contacts are somewhat base-agnostic (Fig. 4, row 2). DBS18 (Fig. 4a, row 2; Fig. 3, row 5) exhibits considerable specificity between the second and 8th position of the HSTelo site; the interactions contributing to this specificity are shown in the zoom in of positions 4-8 (Fig. 4b, row2). Low specificity at position 6 is also consistent with the lack of contact at that position (Fig. 4b, row 2). There is also little to no specificity for the last four positions (Fig. 4c, row 2), because the extended flexible loopy protein finger in this region (Fig. 4a, row 2), provides no major groove base contacts (Fig. S15b, row 2) and is likely flexible given the lower average AF3 pLDDT of the interacting protein residues in the region (Fig. 4c, row 2). DBS20 against the same target (Fig. S13) lacks the loop contacts and has slightly lower affinity and some interaction with one off-target (Fig. S13), but is more specific in terms of the effect of single mutations (Fig. 4, row 1). This design achieves very high specificity for the central GTTA sequence, consistent with the dense network of sidechain-base interactions involving both DNA strands in this region (Fig. 4b, row 1). T5 is the most specific position consistent with the bidentate recognition of the complementary A by Asn49. Selectivity is more modest but sustained over the full 12 base pair (bp) target site, consistent with the interactions with one of the two strands over the full site; only 4 of the 36 single site variants have higher binding affinity than the original target.

DBS22 (Fig. 3, row 6) targets the binding site of the NF*ω*B (Table 1). Unlike the native TF (PDB ID:1NFK) this design is monomeric and has two novel helical sub-domains enabling extensive base contacts. Interface i showcases Arg43 interacting with a dinucleotide G15-G16 step Arg 95 and Thr92 jointly recognizing T7 and G17. Interface ii showcases bidentate recognition of A5 via Gln90, as part of an extensive buttressing network of polar sidechains and the DNA backbone, and base contact with A18 and T6 by Arg95 and Ser91. Together, these interactions likely contribute to the high affinity (Fig. 3c, row 6) and on-target selectivity (Fig. 3d, row 6) of this design. Variant competition assay for DBS22 show strong specificity towards the central four positions (ATTC) of the target (Fig. S18).

Design DBS29 (Fig. 3, row 7) targets a homeodomain target site (HD-site) (Table 1). One of the two sub-domains of DBS29 can be viewed as an extension of the homeodomain fold, with three supporting helices instead of 2 (Fig. 3a, row 7). Asn140 and Gln143 in DBS29 make bidentate interactions with bases A6 and A18, respectively. DBS29 makes extensive minor-groove contacts (Fig. 3a,b, interface i) with the narrow minor-groove of the AT rich core ^15,34,37,38^. In interface ii DBS29 makes extensive hydrogen bonding with 5 consecutive phosphate moieties, indicating indirect recognition of the bound DNA conformation of this flexible target, which lead to specific base recognition as evidenced by the competition assay (Fig. 4, row 4). The BLI experiments indicate a strong affinity of 13 nM (Fig. 3, row 7), and the variant competition experiments show strong on-target selectivity (Fig. 3d, row 7). DBS29 (Fig. 4a, row 5) has high specificity (Fig. 4c, row 5) despite lower average AF3 pLDDT and major groove Hbonds (only two positions 6 and 7 are recognized by major groove bidentate interaction (Fig. 4b, row 5; Fig. 3b, row 7). This suggests an indirect readout mechanism via backbone hydrogen bonds and minor-groove contacts (Fig. 3b, row 7) recognizing the distinct shape ^39^ and flexibility of the AT-rich target^40^.

The majority of these designs have at least seven positions with base-specific interactions as assessed by single mutant variants demonstrating considerable fine-grained specificity. The variation in specificity at different positions could reflect either differences in the extent of protein-DNA contacts, or lack of rigidity of the protein structure in the region. To probe the former, we carried out *in silico* specificity prediction by NA-MPNN ^8^ and DeepPBS ^9^, which considers only the folded structure. These results broadly corroborate the observed specificity for individual base changes for 4 of the 5 designs in Fig. 4, including some of the off-target effects (e.g. off-target preference towards C for DBS20 at position 9 (Fig. S16)). We also investigated if AF3 minPAE is indicative of variant selectivity but we found no clear signal (Fig. S17). To probe the effect of lack of rigidity, we considered the average per residue AF3 confidence (pLDDT) of the contacting protein sidechains per base pair position (upper row in Fig. 4c). These values are generally relatively lower for loopy protein regions and/or regions with less protein-DNA contacts, showing correlation with non-specificity to some extent (e.g. DBS14 positions 8-10, DBS29 positions 1-4, DBS5 positions 4 and 12, DBS18 positions 9-12). Overall, AF3 pLDDT appears to be broadly predictive of information content, requiring only sequence input; NA-MPNN and DeepPBS were predictive of both information content and some variant effects, but requiring a well-folded model as input. Additional designs for which affinity was quantified Fig. S13, all showed low nanomolar affinity. Additional competition assay data are presented in Fig. S18 and curated HT-SELEX specificity profiles for designs (e.g., DBB27, targeting Oct4-gRNA1, which show a binding motif of length 6 with 4 positions matching the intended target) are presented in supplementary information.

## Discussion

Achieving sequence specific DNA recognition is substantially more challenging for *de novo* protein design than protein-protein specificity, as DNA sequences share similar global structures, in contrast to the diverse shapes of protein targets. Consequently, “on-target” positive designs, which are often sufficient to yield specific protein binders without explicit negative design, proved inadequate for DNA binders: designs optimized solely for binding affinity typically exhibited broad, non-specific interactions. We addressed this challenge by integrating multiple complementary strategies. First, we leveraged the power of generative diffusion methods to create proteins that engage with the DNA over an extended region, enabling numerous base-specific contacts. Second, we applied explicit negative design ^41^ to eliminate candidates predicted to bind a set of off-target sequences. Third, we prioritized designs with extensive hydrogen-bonding networks, both to DNA bases for direct readout and to the phosphate backbone to support indirect readout of subtle differences in shape ^15^. We show that RFD3, coupled with explicit *in silico* specificity screening using AF3, can generate sequence selective binders across diverse DNA targets. Modifications to the design pipeline (see supplementary information) enable targeting DNA with non-standard conformation (Fig. S19), further extending our approach. The success rate of obtaining specific designs is nearly two orders of magnitude higher than in previous *de novo* DNA binder design efforts ^12^, which relied on extensive experimental screening to identify sequence selective binders. Our approach enables the generation of DNA binding proteins for a broad range of biologically and therapeutically relevant targets, opening the door to a new class of *de novo*–designed transcription factors and genome-modifying proteins. An important next step will be to characterize target specificity in cellular contexts using ChIP-seq ^42,43,44^ and related methods. Achieving specificity at the level of individual genomic loci will likely require recognition of longer DNA sequences, for example through modular combinations of designed domains, either via rigid covalent fusions ^12^ or engineered heterodimeric interfaces ^12,45,46^. It is informative to compare the architectures of the designed binders to those observed in nature. Structural similarity searches reveal no close global matches between our designs and protein–DNA complexes in the PDB, indicating that the generated folds are largely novel (Fig. S11) while having binding affinities comparable to natives ^47^. At the same time, the designs share features with natural transcription factors, including extended protein–DNA interfaces and clusters of base-contacting hydrogen bonds reminiscent of homing endonucleases ^48,49,50^. Such enzymes achieve high specificity by recognizing longer DNA sequences (typically ~20 bp, compared to ~7–9 bp here), suggesting that increased interface length could further enhance selectivity. This could be achieved either by combining designs generated using the current approach or by extending chain lengths during RFD3-based design. *In silico* tests with longer lengths of proteins indicate reasonable AF3 pass rates (Fig. S20) and higher protein pLDDT in complex with DNA (Fig. S20), suggesting sampling this space could be interesting. Structurally, some designs adopt modular helical bundle architectures reminiscent of homeodomain proteins connected by flexible linkers ^51,52,53^, whereas others form more integrated, monolithic folds distinct from most natural transcription factors ^54,55,56,57^. The structural diversity observed in Fig. 3 and Fig. 4 highlights the ability of generative diffusion models to explore a broad solution space, extending beyond architectures sampled by natural evolution, complementing current family specific computational methods ^3^ for designing DNA binders. While the ability to design sequence specific DNA binding proteins is a substantial advance for *de novo* protein design, there is still considerable room for improvement in design methodology. For approximately half of the targets, no sequence selective binders were identified in sets of 96 designs, and specificity was not absolute even within the limited set of off-targets tested. Moreover, the observed decrease in expression with increasing stringency of specificity selection suggests that scaling to genome-wide specificity may present additional challenges. Several methodological improvements could address these limitations. First, rather than optimizing for binding followed by *in silico* specificity filtering, it may be possible to directly optimize for specificity using inference time search ^58^, enabling more balanced trade-offs between folding, binding, and specificity. Second, incorporating orthogonal structure prediction methods during iterative design and final evaluation (as in ProteinHunter ^21^ and HalluDesign ^22^) could reduce overfitting and improve experimental success rates. With continued advances in design methodology, combined with strategies to extend binding interfaces through multimerization or domain fusion, targeting specific genomic loci with designed proteins will become possible in the near future.

## Data and code availability

The RFD3 checkpoint used in this work is available at https://files.ipd.uw.edu/pub/dna_binder_rfd3/rfd3-1030-foundry.ckpt; Summary metrics of the designs presented can be found at: https://files.ipd.uw.edu/pub/dna_binder_rfd3/summary_data.csv. Further requests for data can be directed to corresponding authors.

## Acknowledgements

We thank Wei Chen, Kate Crawford, Han Raut Altae-Tran, Stephanie Sansbury, Pooja Bandwane, Eric Sun, Zihao Song, Yanjing Li and Rafael Brent from the Institute for Protein Design; Trevor Siggers from Boston University; David Ross, and James McLellan from National Institute of Standards and Technology; and Remo Rohs from University of Southern California for helpful discussions. We thank Kandise VanWormer and Luki Goldschmidt for maintaining the wet lab and computational resources at the Institute for Protein Design. This work was funded by National Science Foundation Graduate Research Fellowship DGE-2140004 (E.S.), Gates Foundation grant INV-043758 and INV-092109 (R.M., R.K., J.B.), Breakthrough Fund Catalyst Design (R.Q.), Fulbright PhD Research grant from the Spanish Fulbright Commission and Spanish State Research Agency grant PID2023-146110NB-I00 (N.G.R.), and the Howard Hughes Medical Institute (D.B.). This research was funded in part by the Wellcome Trust [grant number 220540/Z/20/A] (J.T.). For the purpose of Open Access, the author has applied a CC BY 4.0 copyright licence to any Author Accepted Manuscript version arising from this submission.

## Author Contributions

Research design: D.B., R.M., Y.P., E.S., C.J.G.; Design pipeline development: E.S., Y.P., R.M., N.G.R., A.K., P.T.K., C.J.G., R.J.P.; Design of DBPs: E.S., Y.P., R.M., N.G.R., A.M.; Experimental validation with yeast surface display: Y.P., E.S., R.M. — Variant competition assay: Y.P. — Affinity quantification: E.S., N.G.R. — HT-SELEX: T.W., R.R., J.T.; RFD3 development and support: R.M., J.B., R.K., P.T.K.; Other model development: P.T.K.; Data analysis: E.S., R.M., Y.P., P.T.K., N.G.R., T.W.; Experimental support: B.B., N.R., I.G., D.K.V., Z.S., P.K.; Supervision and organization: D.B., C.J.G., R.M., Y.P., E.S.; Code contribution: E.S., R.M., Y.P., P.T.K., N.G.R., A.K.; Writing: R.M., E.S., Y.P., P.T.K., N.G.R., T.W.; Figures: E.S., R.M., Y.P., P.T.K., N.G.R., — review and editing: All authors.

## Methods

### *De novo* design pipeline of DNA-binding proteins

The *de novo* DNA binder design pipeline is made up of two pipeline segments: the binder block and the specificity block. For the binder block, DNA targets were chosen and generated using AF3 ^20^ (see Target selection and preparation). Protein backbones were diffused for each target (see RFdiffusion3), followed by sequence design using LigandMPNN ^19^ (see LigandMPNN sequence design). Protein sequences were folded with their on-target DNA sequences using AF3 (see Structure prediction). AF3 predictions of less than 8Å DNA-aligned protein C*ε*-RMSD were taken through a resampling round of LigandMPNN sequence design. Resampled sequences were folded using AF3. Predictions less than 3Å DNA-aligned protein C*ε*-RMSD, greater than 0.7 ipTM, and numerous hydrogen bond counts were taken as passing designs from the end of the binder block (Fig. S1).

To enter the specificity block, passing designs from the binder block were filtered with minPAE less than 1.25. These designs were taken through a resampling round of LigandMPNN sequence design (100 sequences per backbone). Protein sequences were folded with their on-target DNA sequences and passing designs were less than 1.5 Å DNA-aligned protein C*ε*-RMSD and greater than 0.9 ipTM. Passing designs were taken into a templated all-by-all AF3 folding where each design sequence was folded in complex with its on-target and six core target DNA sequences (and ten additional target DNA sequences if the on-target for a design was in the additional target set). Δ*minPAE* was computed for each design (see Filtering Metrics) and designs were ranked in decreasing order of this metric. The top 96 designs for each DNA target were ordered as a synthetic oligo for further wet-lab testing (Fig. S1).

### Target selection and preparation

A core set of six therapeutically relevant DNA target sequences was selected to benchmark the design pipeline. Targets were chosen such that no unique 4-mer (or its reverse complement) was shared between sequences, minimizing unintended overlap in binding motifs. The six targets are: GGTGAAATGA (Oct4-gRNA1), GGGCTTGCGA (Oct4-gRNA2), CAGCAGCAGCAG (CAG), ACCTTCTAGG (PCSK9-gRNA), TGAGGAGAGGAG (PRNP-site), CGTATAAACG (TBP-site). To further evaluate generalizability, an additional set of ten targets was selected without k-mer overlap constraints. These sequences were derived from high–information content position weight matrices (PWMs) in the JASPAR database ^60^ and represent diverse transcription factor binding motifs. The ten targets are: ACCACGTGGT (Ebox), GCTTAATTAGCG (HD-site), AGACATGTCT (P53-site), AGGTGTGAAG (Tbox), GGGGATTCCCCC (NFκB-site), GCGTAAACAA (FKH-site), TGACCTTTGA (C4ZF-site), AGGGTTAGGGTT (HSTelo), TCATCCAGCAG (Dux4-gRNA1), GGGCTTGCGA (Dux4-gRNA2). We took the monomeric information-rich core of these entries (Table 1), and padded with a random combination of GC to make the total length 10 or 12 base pairs, whichever was closest. Each target sequence was folded into a double helix structure using AF3^20^ (seed 42; single diffusion sample) to be used as the starting DNA structure for the design pipeline.

### RFdiffusion3

RFdiffusion3 (RFD3) ^18^ was used to generate *de novo* protein scaffolds conditioned on-target DNA structures generated using AF3^20^. For each target DNA structure, scaffolds were generated using multiple ori tokens (center of mass conditioning as described in RFD3^18^), one per six consecutive base-pairs. For a given 6 base-pair stretch of the double helix the ori token is placed at a distance of 3Å towards the major groove, from the centroid of the given stretch, perpendicular to the helical axis (Fig. S1). For each sampled ori, 5100 protein scaffolds (of length in the range 120-150) were generated. Hydrogen bond conditioning ^18^ was applied during generation on candidate ^65^ major groove donor and acceptor atoms (Fig. S1). The is_non_loopy feature was set to True. Target DNA was kept fixed during diffusion. This choice was informed by the monomeric DNA binder generation benchmark of RFD3 ^18^. Model weights used are from the same training trajectory as released in RFD3 ^18^ but trained for longer (see Data and Code availability).

### LigandMPNN sequence design

For every generated scaffold 5 LigandMPNN ^19^ sequences were sampled under a sampling temperature of 0.1. The input complex (the diffusion generated protein backbone) was relaxed using Rosetta FastRelax ^66^ before LigandMPNN sampling.

### Structure prediction

AlphaFold3 (AF3) ^20^ was used to predict protein-DNA complexes from protein sequences generated by LigandMPNN ^19^. Templates were not used throughout the design campaign with the exception of the all-by-all folding step in the specificity block (Fig. S1). For templating, the most recent AF3 prediction before the all-by-all folding was used as the template for the protein chain.

### LigandMPNN sequence design (resampling)

In the binder block, AF3 predictions of designs with less than 8Å DNA-aligned protein C*ε* RMSD were chosen to take into a resampling step (Fig. S1). For each design, 5 LigandMPNN ^19^ sequences were created using a sampling temperature of 0.1. In the specificity block, AF3 predictions of passing binder block designs with minPAE less than 1.25 were resampled with LigandMPNN creating 100 sequences per backbone using a sampling temperature of 0.1 (Fig. S1).

### Filtering metrics

At each AF3 prediction step during the computational pipeline (Fig. S1) a global protein C*ε*-Root Mean Square Deviation (C*ε*-RMSD) when aligned on DNA structure ^18^, predicted interface TM score (ipTM) and minimum Predicted Aligned Error (minPAE) ^20^ was computed between the design model and the AF3 prediction of the protein-DNA complex. Protein-DNA interaction metrics for quantifying generated designs were computed using DSSR ^59^ (v1.7.8). Total hydrogen bonds are quantified as count of all hydrogen bonds returned by DSSR and annotated as ‘nt:aa’. Major groove hydrogen bonds are determined by subsetting to the nucleic acid atoms exposed in the major groove edge of DNA bases. Supporting hydrogen bonds are defined as protein-protein (aa:aa) hydrogen bonds (except when involving protein backbone) to protein residues involved in hydrogen bonds with DNA. For specificity filtering, !*minPAE* metric was calculated by taking the minimum value of the off-target minPAE values and subtracting the on-target minPAE.

Let, P be the set of protein residues and D be the set of DNA residues. We defined minPAE for protein-DNA pairs as,

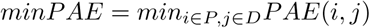

Thus Δ*minPAE* can be defined as,

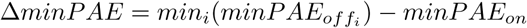

Designs were ranked by their Δ*minPAE* in decreasing order, and the top 96 designs were ordered per DNA target (Fig. S1, Fig. S4).

### Pipeline Ablations

We compared the *in silico* performance of the design strategy when different protein backbone structure generation methods were used to generate initial binder backbones. We compared CA-RFdiffusion ^23^, RFdiffusion All-Atom ^24^, Boltzgen ^25^ and RFD3 ^18^ to generate initial binder backbones. CA-RFdiffusion and RFdiffusion All-Atom were fine-tuned for the task of making DNA binding proteins (Fig. S21). For this purpose we ran a mini-design campaign on 6 target DNA sequences, with 1000 backbones generated per target with each method, 5 ligandMPNN ^19^ sequences generated per backbone, then folding these sequences with AF3^20^. Pass rates of designs were calculated using DNA-aligned protein RMSD (Fig. 1). Biophysical metrics of passing designs were also computed as described in the Filtering Metrics section (see Filtering Metrics).

### Evo-1 embeddings of TF sequence space

To map vertebrate transcription factor (TF) binding sequence space, we obtained position frequency matrices (PFMs) for all vertebrate TF motifs from the JASPAR database ^60^ via the REST API. The top 10 TFs per family ranked by information content were selected, with 5 sequences sampled per TF, yielding 2,355 sequences across 91 TF families. Each sequence was embedded using Evo-1-131k-base, a 7-billion parameter genomic language model ^67^. Sequences were passed individually (no padding) through the model, and the final-layer hidden states were mean-pooled across sequence positions to produce a single 4,096-dimensional embedding vector per sequence. Dimensionality was then reduced to two dimensions using PCA fit exclusively on the TF sequence embeddings. Designed non-native DNA-binding protein target sequences were embedded in the same Evo session on the same GPU and projected into the TF PCA space using the fitted transform, without influencing the PCA axes.

### ESM-2 sequence embedding and dimensionality reduction

Protein sequences were extracted from crystal structures of protein–DNA complexes deposited in the Protein Data Bank ^29,30^ (n = 532 unique chains). For each structure, protein chains were identified by filtering for standard amino acid residues with a minimum length of 20 residues. *De novo* designed DNA-binding protein sequences (n = 73) spanning 9 target sites were extracted from AlphaFold3-predicted structures. Per-protein representations were obtained using ESM-2^68^ (650M parameters), a protein language model pretrained on approximately 250 million sequences from UniRef50. Each sequence was passed through the full 33-layer transformer, and the beginning-of-sequence (BOS) token representation from the final layer (1,280 dimensions) was used as the whole-protein embedding. Because the BOS token attends to all residue positions through successive self-attention layers, its final-layer representation encodes a global summary of the sequence without requiring pooling across variable-length outputs. Principal component analysis (PCA) was fitted on the native crystal structure embeddings, and designed sequences were projected onto the same principal component axes.

### DNA library preparation

For high-throughput screening of the binder block order, protein sequences were padded to 128 amino acids by appending a (GGS)*_n_* linker at the C terminus to satisfy the length constraint (<15% variation) required for pooled synthesis on a 500nt gene fragment chip from Twist Bioscience. Sequences were reverse translated and codon optimized for S. cerevisiae using DNAWorks ^69^. The resulting oligonucleotide library was synthesized as a pooled 500-nt oligo pool. Lyophilized oligo pools were resuspended in nuclease-free water to 100 ng *µ*L^⎯^1 and stored at -20 °C. Prior to amplification, oligos were diluted to working concentrations for quantitative PCR. The 5’ and 3’ library fragments were independently amplified using custom primers and KAPA HiFi HotStart polymerase, with reactions monitored by qPCR and terminated in mid-exponential phase to minimize amplification bias. PCR products were purified between steps by gel extraction or column purification. Internal primer sequences were removed using USER enzyme treatment followed by end repair according to manufacturer protocols. Processed 5’ and 3’ fragments were assembled by overlap PCR under the same amplification conditions. Assembly products were verified by agarose gel electrophoresis, and bands of the expected size were excised and purified. Purified libraries were re-amplified to obtain sufficient material for yeast transformation. For each transformation, 2–6 *µ*g of insert DNA was combined with linearized pETCON3 vector at an approximate 3:1 molar ratio and electroporated into electrocompetent S. cerevisiae EBY100 using a previously described protocol ^70^. For deep sequencing, libraries were amplified from plasmid DNA extracted from sorted yeast populations. Plasmid extraction was performed using Solution 1 Digestion Buffer from Zymo Research. Illumina sequencing adapters and sample-specific barcodes were introduced during a second PCR step, followed by gel purification. Sequencing was performed on an Illumina NextSeq platform (Illumina).

### Yeast surface display

Yeast surface display was performed using the S. cerevisiae EBY100 strain. Cultures were grown in C–Trp–Ura medium supplemented with 2% (w/v) glucose (CTUG). For induction, CTUG cultures were diluted 1:10 into SGCAA medium containing 0.2% (w/v) glucose to a final density of 1*×*10*^-7^* cells mL^−^1 and incubated at 30°C for 16–24 h. Induced cells were washed with PBSF (PBS supplemented with 1% (w/v) BSA) and labeled with 5’BiotinTEG-biotinylated double-stranded DNA targets using either an avidity or non-avidity staining protocol. In the avidity protocol, cells were incubated simultaneously with biotinylated DNA target, anti-c-Myc FITC antibody (Miltenyi Biotec), and streptavidin–phycoerythrin (SAPE; Thermo Fisher Scientific). SAPE was used at one quarter the molar concentration of the biotinylated DNA target. In the non-avidity protocol, cells were first incubated with a biotinylated DNA target, washed, and subsequently labeled with SAPE and FITC. After labeling, cells were washed in PBSF prior to analysis or sorting. Cell sorting of labeled yeast libraries (for the binder block) was performed on a Sony SH800S cell sorter (Sony Biotechnology). Libraries were first sorted using the avidity protocol at 1 *µ*M target concentration to remove weak binders, followed by an enrichment sort at 1 *µ*M using the non-avidity protocol. Subsequent titration sorts were performed at decreasing target concentrations (100 nM, 10 nM, 1 nM, 0.1 nM) to enrich for higher-affinity binders within the pools. To test for on-target binding activity of the binder block designs, coding sequences were synthesized as gene fragments (Twist Bioscience), transformed into yeast and induced in 96-well plate format. Labeling with 1*µ*M of respective on-target biotinylated DNA targets and SAPE/FITC was performed in plates and analyzed by flow cytometry using an Attune N×T with autosampler (Thermo Fisher Scientific). Flow cytometry data were analyzed using custom Python scripts and the CytoFlow package. For each sample, the expressing population was gated using a Gaussian mixture model implemented in CytoFlow, and the ratio of SAPE fluorescence to FITC fluorescence (binding signal normalized to expression) was calculated for all gated events. For EC_50_ measurements, individual designs were subjected to an 11-point titration binding assay (3 *µ*M to 0.152 nM) of its on-target biotinylated dsDNA target using the same labeling and analysis workflow. The binding-to-expression ratio at each concentration was fit using a Hill equation to estimate EC_50_ values. To assess binding specificity, the yeast display workflow and analysis were extended to an all-by-all binding assay. Each design was tested at 1*µ*M concentration against the full panel of biotinylated dsDNA targets, with each protein–DNA pair assayed in a separate well. Background signal was defined using no-target control samples and subtracted from all measurements. The binding-to-expression signal ratio for each condition was compiled and visualized as a heatmap to quantify target specificity across the library. Yeast surface display-based competition assays were performed as previously described ^12^. Cells were incubated under non-avidity conditions with 1 *µ*M 5’ BiotinTEG dsDNA duplex oligonucleotides in the presence of 8*µ*M non-biotinylated competitor dsDNA duplex oligonucleotides. Cells were screened and analyzed using the yeast display workflow described above. For each sample, median PE/FITC values were calculated. Background signal was defined using no-target control samples and subtracted from all measurements. Data were subsequently normalized to the median signal from no-competitor control samples to enable comparison across conditions.

### Generation of yeast displaying individual protein designs

To characterize designs from the specificity block, DNA fragments for the expression and screening of protein designs (IPD blocks) were generated from oligonucleotide pools, similar to previous reports ^71,72^. Briefly, DNA sequences encoding designed proteins were bioinformatically fragmented, and each fragment was appended with BsaI restriction sites and design-specific PCR priming sites. These sequences were synthesized within a 350 nucleotide oligonucleotide pool at Twist Bioscience. Two nested PCR stages were conducted to dial-out design-specific fragments within each well of a microwell plate. DNA fragments for each design were individually combined with a backbone containing pETCON adapter sequences (CF_2) and assembled via Golden Gate reaction. Assembled constructs were amplified into linear DNA fragments via PCR using pETCON-compatible primers with repliQa HiFi ToughMix and EvaGreen dye for real-time monitoring. Unpurified PCR products encoding each design were co-transformed with linearized pETCON3 vector into chemically competent EBY100 S. cerevisiae via lithium acetate heat-shock transformation and gap repair ^73^. Transformations were performed in 20 *µ*L reactions containing 2.5 ng/µL digested pETCON3 and 3 *µ*L PCR insert, with heat shock at 42°C in 45% PEG 3350. Transformants were recovered in CTUG selective media. On-target binding and specificity assays were performed as previously described (see Yeast surface display).

### Deep sequencing analysis

For the analysis of the binder block designs, paired-end reads from yeast sorting experiments were merged using PEAR software ^74^ and sequenced on a NextSeq 2000 platform from Illumina. Merged reads were translated *in silico* and matched to the designed protein library to obtain per-design read counts for each sorted pool. To reduce the impact of sequencing noise, reads corresponding to a given design that represented less than 1×10^-4^% of total reads in a pool were discarded. For each pool, enrichment of a design was calculated as the log_2_ ratio between the fraction of reads for that design in the sorted pool and the fraction of reads for the same design in the naïve expression pool. Designs were considered enriched if the on-target log_2_ enrichment exceeded 1.5. Candidates were selected for follow-up characterization if their enrichment against off-target DNA pools was lower than their enrichment against the intended on-target pool, indicating sequence specificity.

### Protein expression and purification

Synthetic DNA fragments encoding *de novo* dna binder monomers were ordered as codon-optimized gBlocks from Integrated DNA Technologies. Fragments were cloned into the LM627 expression vector by Golden Gate assembly using BsaI overhangs and transformed into BL21(DE3) E. coli. Single transformations were used to inoculate 25mL starter cultures in LB supplemented with 25 mgL*^-1^* kanamycin and grown overnight at 37°C. For expression, 10mL of starter culture was used to inoculate 490mL LB with kanamycin and grown at 37°C to an OD_600_ of 0.6–0.8. Protein expression was induced with IPTG, and cultures were incubated at 18°C for 16–18h. Cells were harvested by centrifugation (3,000g, 15min) and resuspended in lysis buffer (300mM NaCl, 25mM Tris-HCl pH 8.0, 5mM MgCl_2_, 0.1mM CaCl_2_, 0.5 mg mL*^-1^* DNase I, 1mM PMSF). Cells were lysed by sonication (10s on/off pulses, 7.5min total, 80% amplitude), and lysate was clarified by centrifugation (10,000g, 30min). Clarified lysate was applied to a gravity column containing 2mL Ni-NTA resin (stored in 50% ethanol). The resin was equilibrated with high-salt wash buffer (2M NaCl, 25mM Tris-HCl, 30mM imidazole, pH 8.0). After loading the samples, the column was washed with 10 column volumes of high-salt wash buffer followed by 5 column volumes of SNAC buffer (100mM CHES, 100mM acetone oxime, 100mM NaCl, 500mM guanidine hydrochloride, pH 8.6). Columns were then capped and incubated overnight at room temperature in 5 column volumes of SNAC buffer supplemented with 0.2mM NiCl_2_ on an orbital shaker. The following day, flow-through was collected and the resin was washed with an additional 5 column volumes of SNAC buffer. Eluted protein was concentrated to 1mL and loaded on a Superdex 75 Increase 10/300 GL column (Cytiva) equilibrated in gel filtration buffer (100mM NaCl, 25mM Tris-HCl, 0.5mM TCEP, pH 8.0). Fractions corresponding to monomeric species were pooled and concentrated to 200*µ*L. Protein concentrations were determined from absorbance at 280nm using calculated extinction coefficients and expected molecular weights (Beer–Lambert law). Protein identity and purity were confirmed by mass spectrometry.

### BLI

Binding measurements were performed on an Octet R8 (Sartorius) at 24°C and 1200 rpm shaking. Streptavidin-coated biosensors (18-5019) were hydrated for 10 min in BLI buffer (1× PBS supplemented with 0.05% BSA and 0.5% Tween-20). Analyte proteins were diluted from concentrated stocks into BLI buffer immediately prior to measurement. Non-specific protein-sensor interactions were assessed by testing binding to unloaded sensors. 5’-biotinylated dsDNA targets (purchased from IDT) at 100 nM were immobilized onto sensors for 30-45s to reach a 0.1-0.2 nm response followed by a 90s baseline in BLI buffer. Protein association signal was recorded on 2:3 serial dilutions over 240s and dissociation into BLI buffer was recorded for 300 s. The traces from loaded sensors in absence of protein were subtracted to correct for baseline drift. Kinetic parameters and dissociation constants (K*_D_*) were determined by global fitting to a 1:1 binding model across the dilution series using the Octet Analysis Studio software.

### HT-SELEX

Target proteins (DBB/DBS) were expressed and purified as described previously ^75^. Proteins were produced using a pETG20A Gateway expression vector containing an N-terminal Thioredoxin–6× His tag and a C-terminal SBP tag, expressed in E. coli Rosetta pLysS, and stored in 50% glycerol at -20°C. HT-SELEX (high-throughput systematic evolution of ligands by exponential enrichment) was performed by incubating 100–200 ng of a random DNA library (comprising 101 bases of hand-mixed random DNA flanked by priming sites) with 100-200 ng of the proteins ^76^ for 20–30 minutes at 25°C in SELEX buffer (31 mM Tris pH 7.5, 119 mM KCl, 1.5 mM MgCl_2_, 6% glycerol, 100 *µ*M EGTA, 1 mM DTT, 3 *µ*M ZnSO_4_). The experiment was optimised such that the effective salt concentration during TF–DNA interactions was approximately 90 mM. TF–DNA complexes were captured using nickel magnetic beads (28-9799-17, GE Healthcare), followed by washing with 5 mM Tris-HCl pH 7.5, 5 mM EDTA to remove unbound DNA. The TF-DNA complexes were then heat-denatured at 80°C for 10 minutes in elution buffer (0.1% Tween, 10 mM Tris pH 7.8, 1 mM MgCl_2_), after which the enriched DNA was recovered, PCR-amplified, and used as input for subsequent cycles. Cycles 2–4 were performed using the same procedure. Libraries from cycles 1, 3, and 4 were indexed using P5/P7 primers and sequenced on an Element AVITI platform (PE150).

### Motif Mining

Demultiplexed reads were merged with fastp ^77^ requiring a minimum overlap of 5 bp, followed by adapter trimming. Only merged reads corresponding to the expected 101 bp variable region were retained for downstream analysis. Primary motifs were manually curated for each target protein from the HT-SELEX libraries. Positional frequency matrices (PFMs) were generated using Autoseed ^76,78^ (multinomial = 1), with seeds selected based on established criteria ^79^.

