## Supplementary Information for "Generative design of sequence specific DNA binding proteins"

for

### Supplementary Figures

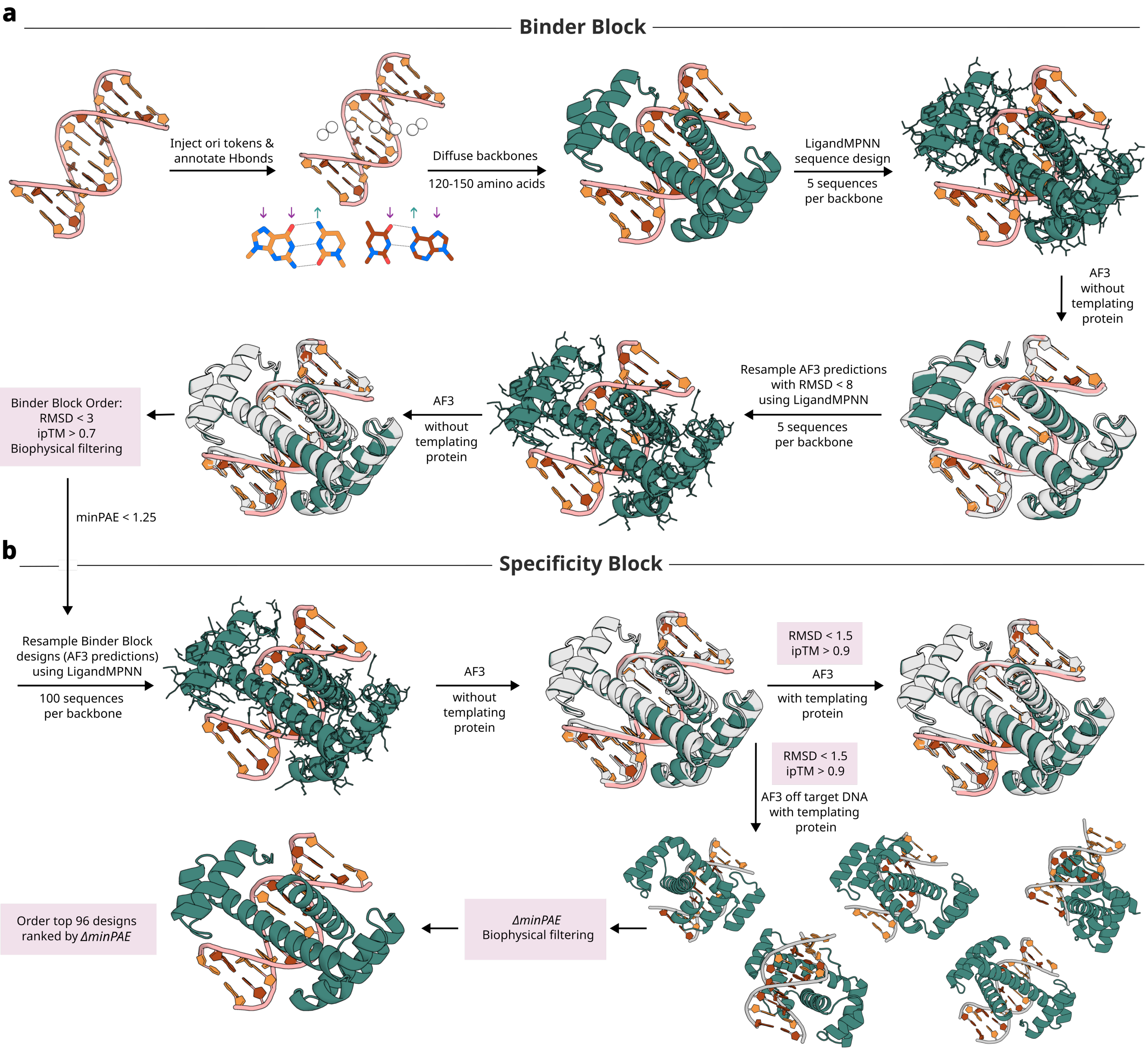
a
Binder Block
Inject ori tokens &
annotate Hbonds
Diffuse backbones
120-150 amino acids
LigandMPNN
sequence design
5 sequences
per backbone
AF3
without
templating
protein
Resample AF3 predictions
with RMSD < 8
using LigandMPNN
5 sequences
per backbone
AF3
without
templating
protein
Binder Block Order:
RMSD < 3
ipTM > 0.7
Biophysical filtering
minPAE < 1.25
b
Specificity Block
Resample Binder Block
designs (AF3 predictions)
using LigandMPNN
100 sequences
per backbone
AF3
without
templating
protein
RMSD < 1.5
ipTM > 0.9
AF3
with templating
protein
RMSD < 1.5
ipTM > 0.9
AF3 off target DNA
with templating
protein
ΔminPAE
Biophysical filtering
Order top 96 designs
ranked by ΔminPAE

**Fig. S1** ***(previous page).*** **Extended depiction of the *de novo* DNA binder design pipeline. a,** Binder block workflow. An initial AF3 prediction is generated for the target DNA sequence. Centers of mass (COMs; white spheres) are defined on the DNA and used as distinct initialization points for diffusion-based backbone generation. Hydrogen-bond donors (green arrows) and acceptors (purple arrows) on the DNA bases are annotated and can be used to guide the diffusion trajectory. Protein backbones (120–150 amino acids) are generated from these configurations using diffusion, followed by sequence design with LigandMPNN. Designed sequences are evaluated with AF3, and candidates with RMSD $< 8$Å are iteratively refined by resampling with LigandMPNN on the AF3-predicted structures. Final designs passing filtering criteria are selected for pooled synthesis and high-throughput screening. **b,** Specificity block workflow. Designs from the binder block with minPAE $< 1.25$ are subjected to additional rounds of sequence optimization with LigandMPNN and AF3 evaluation. Specificity is assessed through all-by-all *in silico* folding, in which each design is modeled with both on-target and off-target DNA sequences using AF3 with the protein structure templated from the latest prediction. Designs meeting structural quality thresholds (RMSD $< 1.5$Å and iPTM $> 0.9$) are further evaluated using $\Delta minPAE$ (see Fig. S4). Candidates are ranked by $\Delta minPAE$, and the top 96 designs are selected for synthesis and plate-scale experimental characterization.

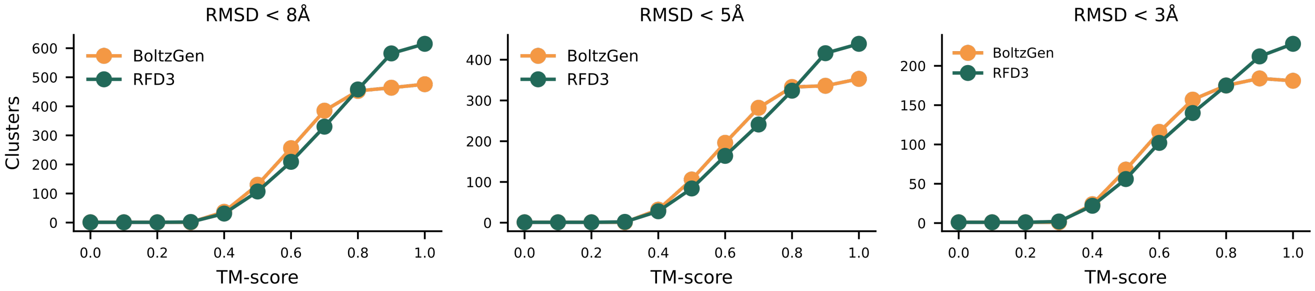

RMSD < 8Å
RMSD < 5Å
RMSD < 3Å
BoltzGen
RFD3
Clusters
TM-score

**Fig. S2** ***(previous page)*****. Structural similarity of designs generated by RFdiffusion3 and BoltzGen.** TM-align[80] scores were computed across generated protein backbones from RFdiffusion3 and BoltzGen (six DNA targets, 1000 backbones per target), and then clustered at various TM-score cutoffs. Backbones were inverse-folded with LigandMPNN and folded with AF3 (5 sequences per backbone). We report the number of backbone clusters with at least one sequence with a refolding RMSD of below 8Å, 5Å, and 3Å at each TM-score cutoff. The models exhibit similar backbone diversity, with BoltzGen sampling a wider space at a lower, more stringent cutoff, but RFdiffusion3 sampling a wider space overall.

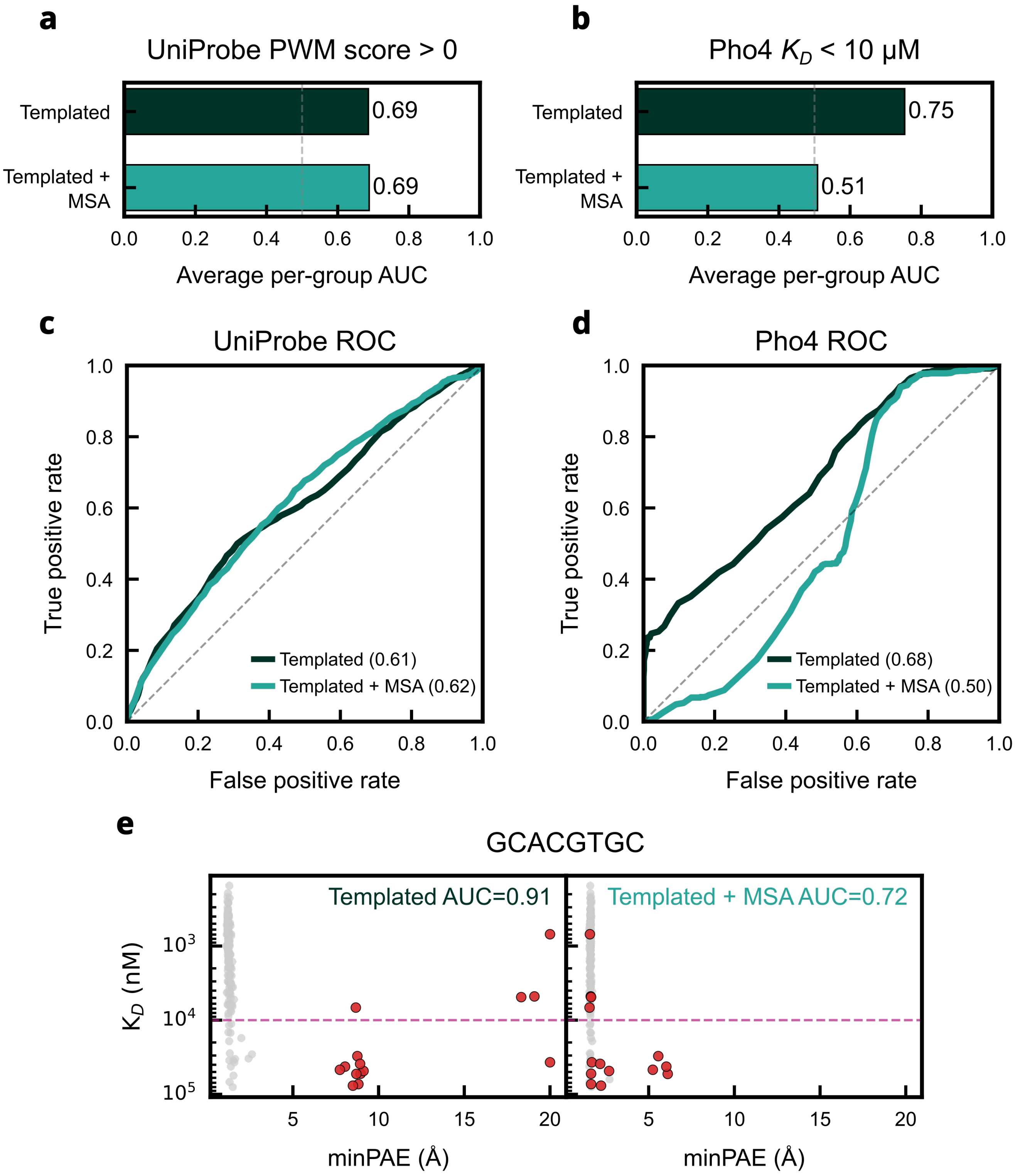
a
UniProbe PWM score > 0
Templated
Templated +
MSA
0.69
0.69
0.0
0.2
0.4
0.6
0.8
1.0
Average per-group AUC
b
Pho4 $K_D$ < 10 μM
Templated
Templated +
MSA
0.75
0.51
Average per-group AUC
c
UniProbe ROC
True positive rate
False positive rate
Templated (0.61)
Templated + MSA (0.62)
d
Pho4 ROC
True positive rate
False positive rate
Templated (0.68)
Templated + MSA (0.50)
e
GCACGTGC
Templated AUC=0.91
Templated + MSA AUC=0.72
$K_D$ (nM)
$10^3$
$10^4$
$10^5$
minPAE (Å)
minPAE (Å)

**Fig. S3** ***(previous page)*****. Templated AF3 minPAE as an indicator of selectivity.** Mean AUC for binding prediction using AF3 minPAE as a predictor, grouped by dataset **a,** UniProbe transcription factors (TFs)[81] **b,** Pho4 TF variants[82]. AUC was computed independently for each group and averaged. A protein–DNA pair was classified as a binder if the PWM score was $> 0$ for UniProbe TFs, and if and if $K_D < 10$ $\mu$M for Pho4 variants. AF3 predictions were performed with and without multiple sequence alignment (MSA) information. Pooled receiver operating characteristic (ROC) curves across all data points for **c,** UniProbe TFs and **d,** Pho4 TF variants; legend values indicate the corresponding pooled AUC. **e,** Scatter plot of binding affinity versus minPAE for the GCACGTGC sequence in the Pho4 dataset, comparing AF3 predictions with and without MSA information. Red points highlight false positives in the +MSA condition (low minPAE values) that are resolved upon removal of MSA information.

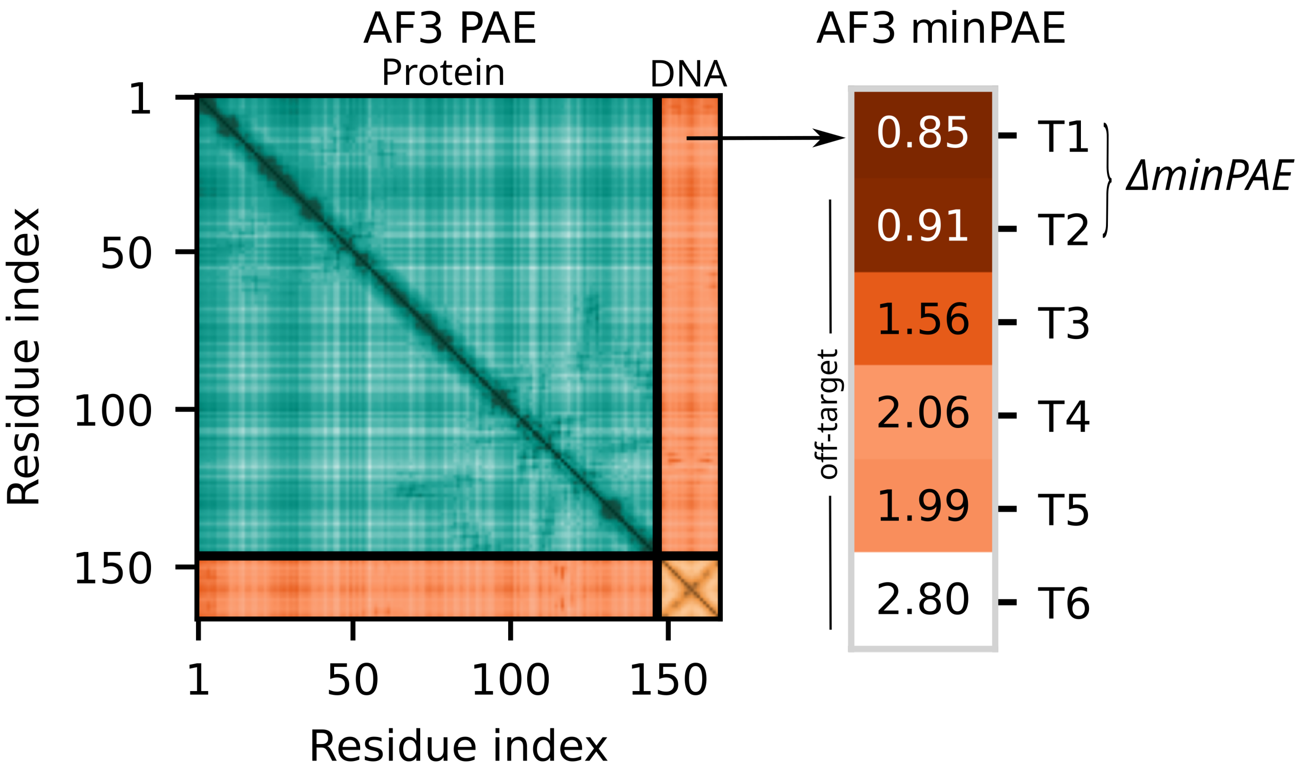
AF3 PAE
Protein
DNA
1
50
100
150
Residue index
1
50
100
150
Residue index
AF3 minPAE
0.85
0.91
1.56
2.06
1.99
2.80
T1
T2
T3
T4
T5
T6
ΔminPAE
off-target

**Fig. S4** ***(previous page).*** **Schematic description of $\Delta minPAE$ calculation.** AF3 predicted aligned error (PAE) values shown as a residue by residue matrix for a predicted complex: protein-protein (teal), DNA-DNA(yellow), protein-DNA(orange). Protein-DNA values are used to derive the minPAE score for a given complex (left side). The resulting minPAE value is computed for a given protein folded across all targets (T1-T6) and the difference between the on-target minPAE (T1) and the best scoring off-target minPAE values (T2-T6) is the $\Delta minPAE$ (see Methods) value for the *de novo* DB under consideration (right side).

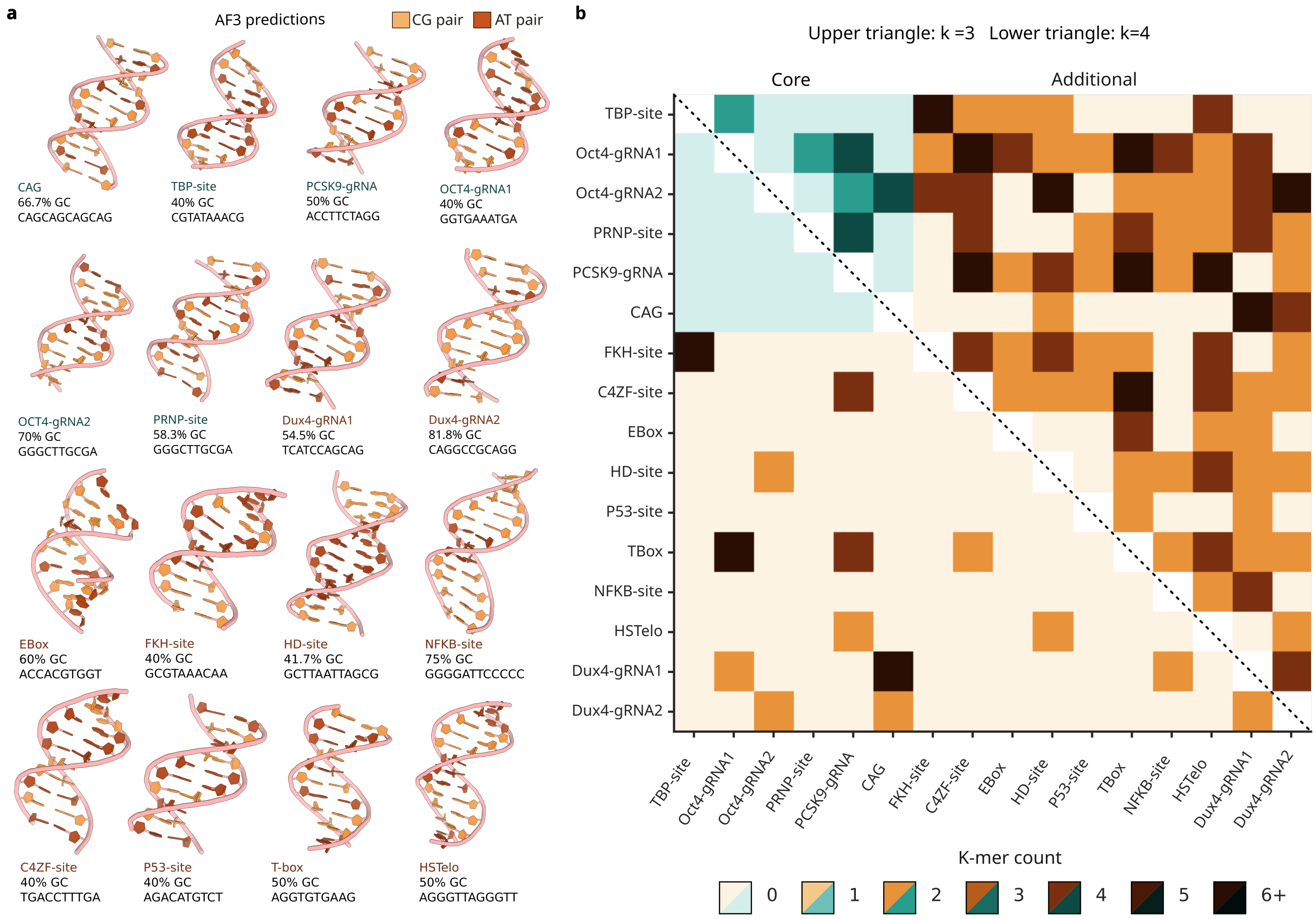
a
AF3 predictions
CG pair
AT pair
CAG
66.7% GC
CAGCAGCAGCAG
TBP-site
40% GC
CGTATAAACG
PCSK9-gRNA
50% GC
ACCTTCTAGG
OCT4-gRNA1
40% GC
GGTGAAATGA
OCT4-gRNA2
70% GC
GGGCTTGCGA
PRNP-site
58.3% GC
GGGCTTGCGA
Dux4-gRNA1
54.5% GC
TCATCCAGCAG
Dux4-gRNA2
81.8% GC
CAGGCCGCAGG
EBox
60% GC
ACCACGTGGT
FKH-site
40% GC
GCGTAAACAA
HD-site
41.7% GC
GCTTAATTAGCG
NFKB-site
75% GC
GGGGATTCCCCC
C4ZF-site
40% GC
TGACCTTTGA
P53-site
40% GC
AGACATGTCT
T-box
50% GC
AGGTGTGAAG
HSTelo
50% GC
AGGGTTAGGGTT
b
Upper triangle: k =3 Lower triangle: k=4
Core
Additional
TBP-site
Oct4-gRNA1
Oct4-gRNA2
PRNP-site
PCSK9-gRNA
CAG
FKH-site
C4ZF-site
EBox
HD-site
P53-site
TBox
NFKB-site
HSTelo
Dux4-gRNA1
Dux4-gRNA2
K-mer count
0
1
2
3
4
5
6+

**Fig. S5** ***(previous page)*****. DNA binding protein target sequences and k-mer similarity analysis.** **a**, AF3 structural predictions of the DNA target sequences, including six core targets (TBP-site, Oct4-gRNA1, Oct4-gRNA2, PRNP-site, PCSK9-gRNA, and CAG) and ten additional targets (FKH-site, C4ZF-site, EBox, HD-site, P53-site, TBox, NFKB-site, HSTelo, Dux4-gRNA1, and Dux4-gRNA2). **b**, Pairwise k-mer overlap matrix across all 16 target sequences, considering both forward and reverse complement strands. The upper triangle shows overlap at k=3 and the lower triangle at k=4. Color intensity reflects the number of shared k-mers (0–6+), with teal indicating overlaps within core targets and orange indicating overlaps involving additional targets. Sequences were designed to minimize shared sequence content, particularly within the core set.

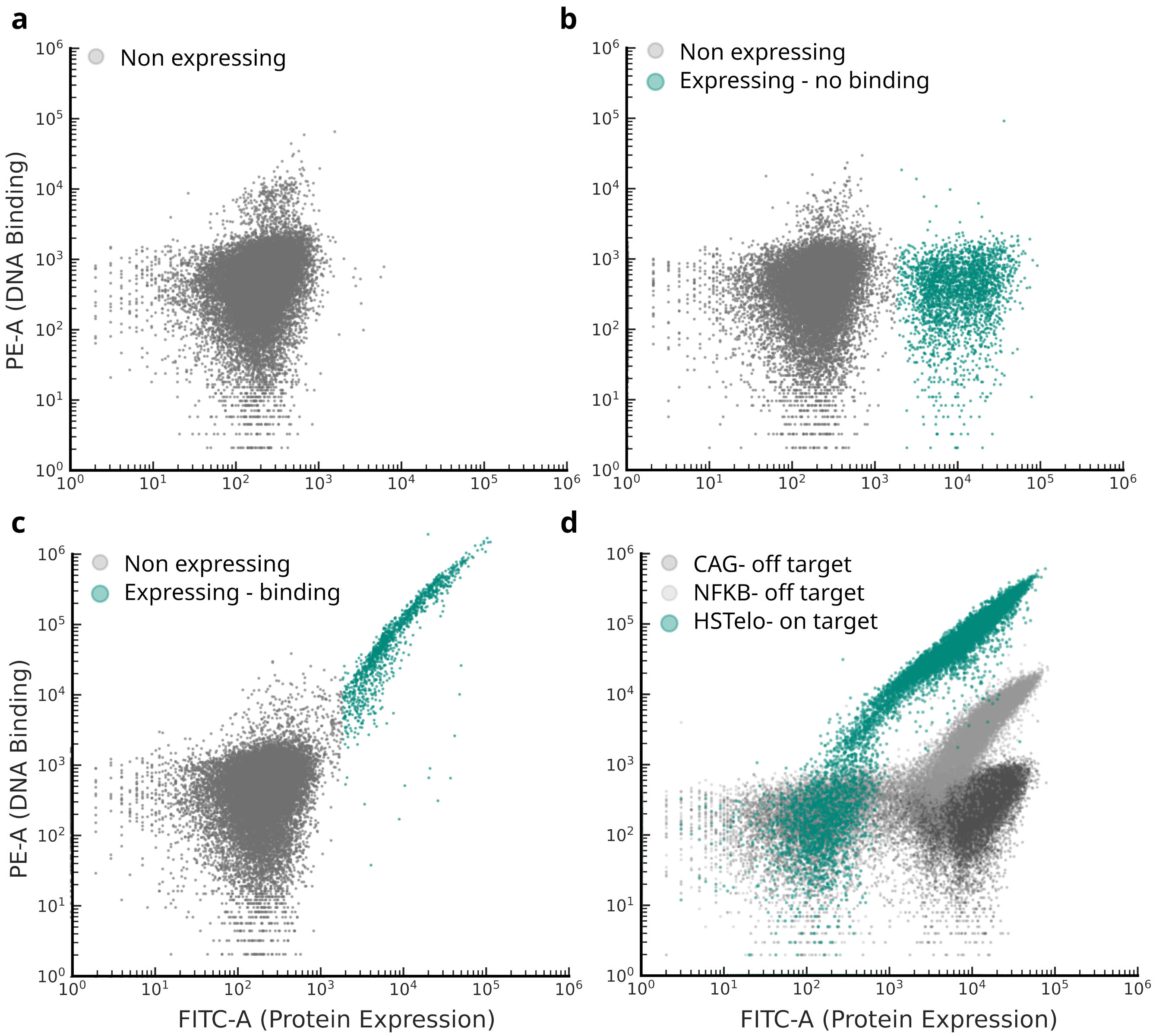
a
Non expressing
b
Non expressing
Expressing - no binding
c
Non expressing
Expressing - binding
d
CAG- off target
NFKB- off target
HSTelo- on target
PE-A (DNA Binding)
FITC-A (Protein Expression)

**Fig. S6** ***(previous page)*****. Representative yeast surface display flow cytometry data.** **a,** Flow cytometry plot showing a non-expressing yeast population ($PE^-/FITC^-$). **b,** Flow cytometry plot showing a non-expressing population (gray; $PE^-/FITC^-$) and an expressing population (teal; $PE^-/FITC^+$). **c,** Flow cytometry plot showing a non-expressing population (gray; $PE^-/FITC^-$) and an expressing, DNA binding population (teal; $PE^+/FITC^+$). **d,** Overlay of representative flow cytometry traces illustrating a range of binding behaviors among expressing populations, including strong binders (teal), intermediate binders (light gray), and weak binders (dark gray).

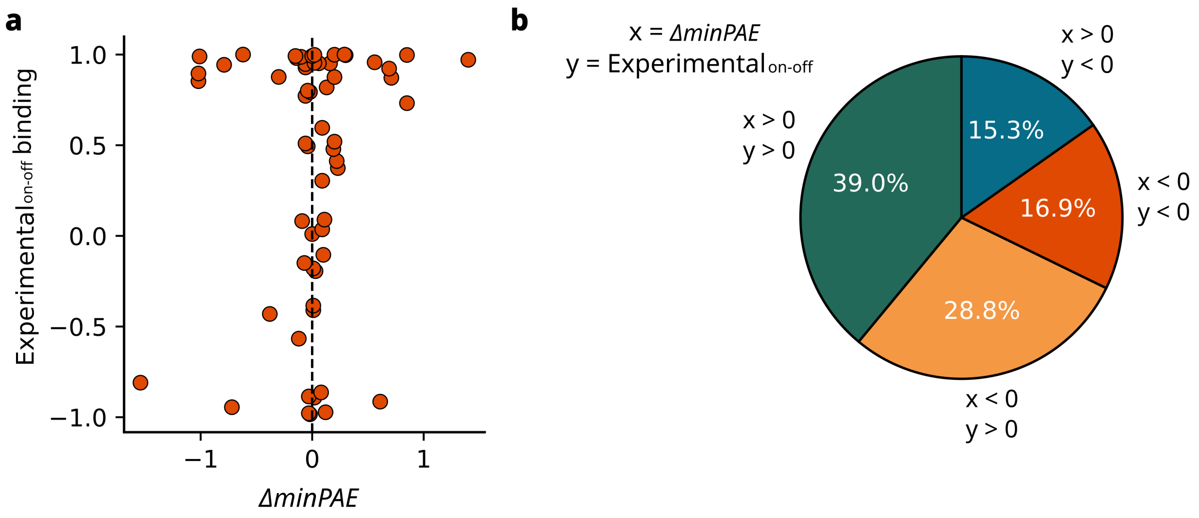
a
1.0
0.5
0.0
−0.5
−1.0
Experimental on-off binding
−1
0
1
ΔminPAE
b
x = ΔminPAE
y = Experimental on-off
x > 0
y < 0
x > 0
y > 0
x < 0
y < 0
x < 0
y > 0
15.3%
39.0%
16.9%
28.8%

**Fig. S7** ***(previous page)*****. Correlation between experimental binding and $\Delta minPAE$. a,** Scatter plot of $\Delta minPAE$ versus experimental binding specificity for individual designs. Experimental specificity is defined as the difference between on-target and strongest off-target normalized binding signal (PE/FITC$_{on}$ - max(PE/FITC$_{off}$)). Analysis was performed on the binder block dataset hits (n = 59). **b,** Distribution of designs by agreement between computational and experimental specificity. Pie chart showing the fraction of designs for which $\Delta minPAE$ and experimental specificity are both positive, both negative, or discordant (opposite signs). Among designs with $\Delta minPAE > 0$ (the applied selection criterion), 71.8% (23 of 32) exhibit experimentally validated on-target selectivity.

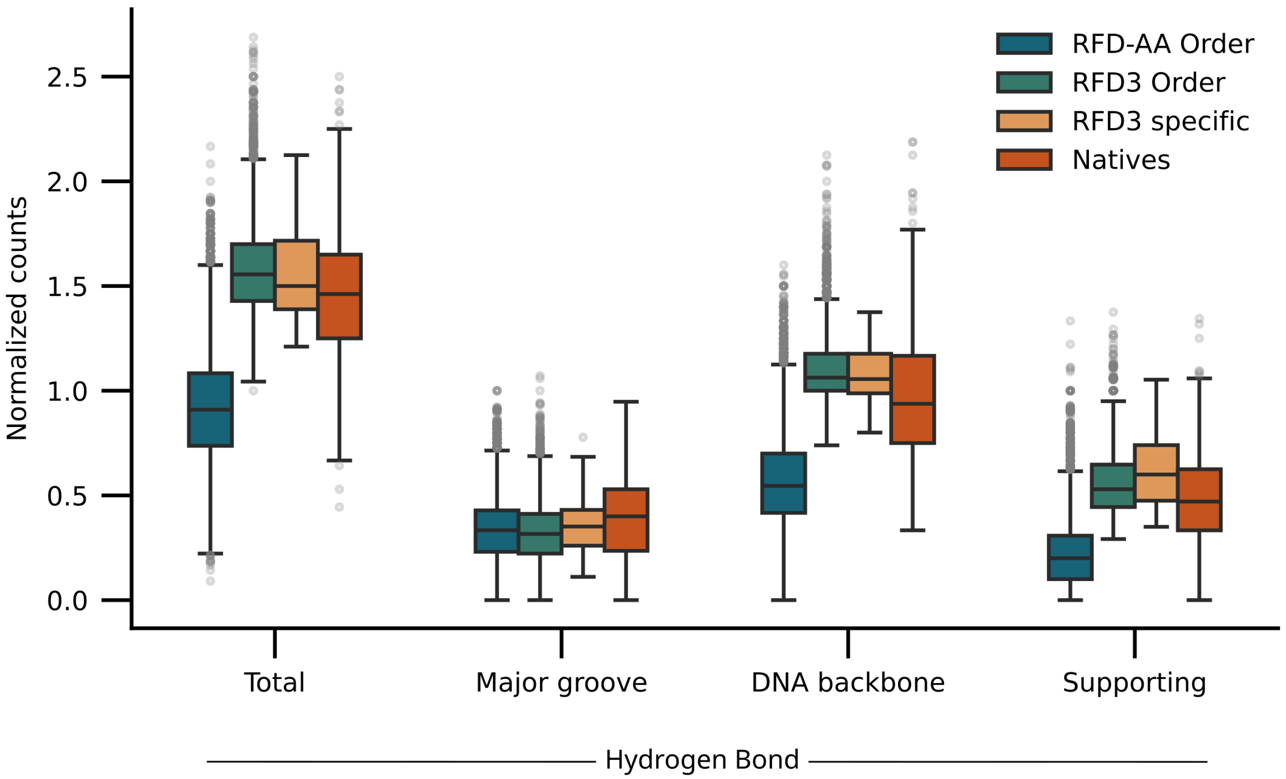

RFD-AA Order
RFD3 Order
RFD3 specific
Natives
Normalized counts
0.0
0.5
1.0
1.5
2.0
2.5
Total
Major groove
DNA backbone
Supporting
Hydrogen Bond

**Fig. S8** ***(previous page)*****. Biophysical comparison of designed and native protein–DNA interfaces.** Per-nucleotide interaction counts for protein–DNA complexes across design campaigns and native transcription factors (TFs). Interactions were computed from AF3 predicted structures using DSSR[59] (v1.7.8). The RFD-AA set comprises 26,575 designs, generated using the RFD-AA pipeline described in (Fig. S21). The RFD3 set comprises 12,653 designs, and the RFD3-specific subset includes 56 designs exhibiting sequence selective binding. Native TFs include 357 structures from the JASPAR database[60] with mean information content >1.5.

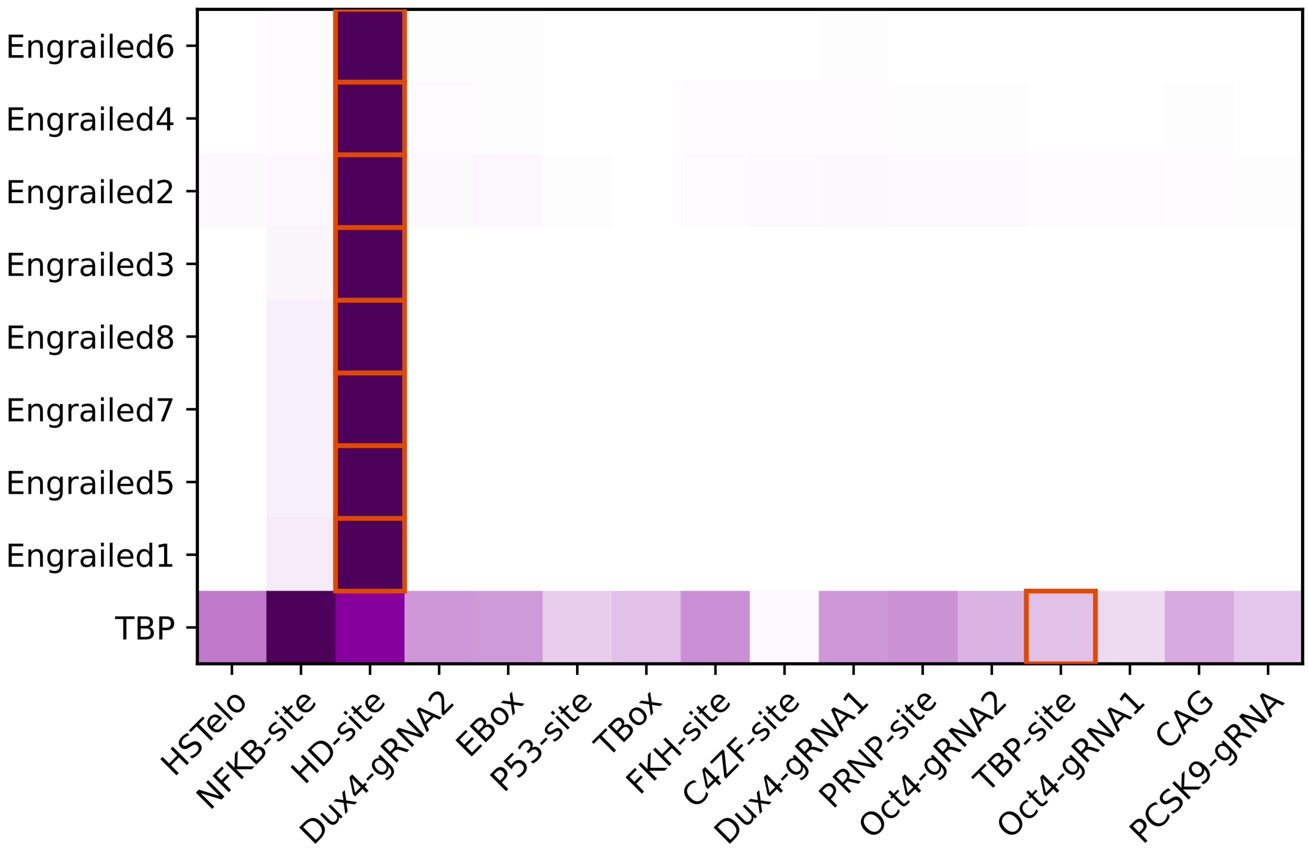
Engrailed6
Engrailed4
Engrailed2
Engrailed3
Engrailed8
Engrailed7
Engrailed5
Engrailed1
TBP
HSTelo
NFKB-site
HD-site
Dux4-gRNA2
EBox
P53-site
TBox
FKH-site
C4ZF-site
Dux4-gRNA1
PRNP-site
Oct4-gRNA2
TBP-site
Oct4-gRNA1
CAG
PCSK9-gRNA

**Fig. S9** ***(previous page)*****. All-by-all specificity of a subset of native DNA binding proteins.** All-by-all specificity matrix for a subset of native TFs across DNA targets, showing PE/FITC binding signal normalized by column. Proteins were screened by yeast surface display at 1$\mu$M DNA (non-avidity format). Red boxes denote on-target sequence for each TF. Unipriot IDs for Engrailed1: P02836, Engrailed2: B4K2E4, Engrailed3: B4MYI0, Engrailed4: B4GAN8, Engrailed5: A0A811VDM2, Engrailed6: A0A7R8U9R3, Engrailed7: A0A8J6H9A6, Engrailed8: A0A6J0BU91, and TBP: A0A7K4JSS6.

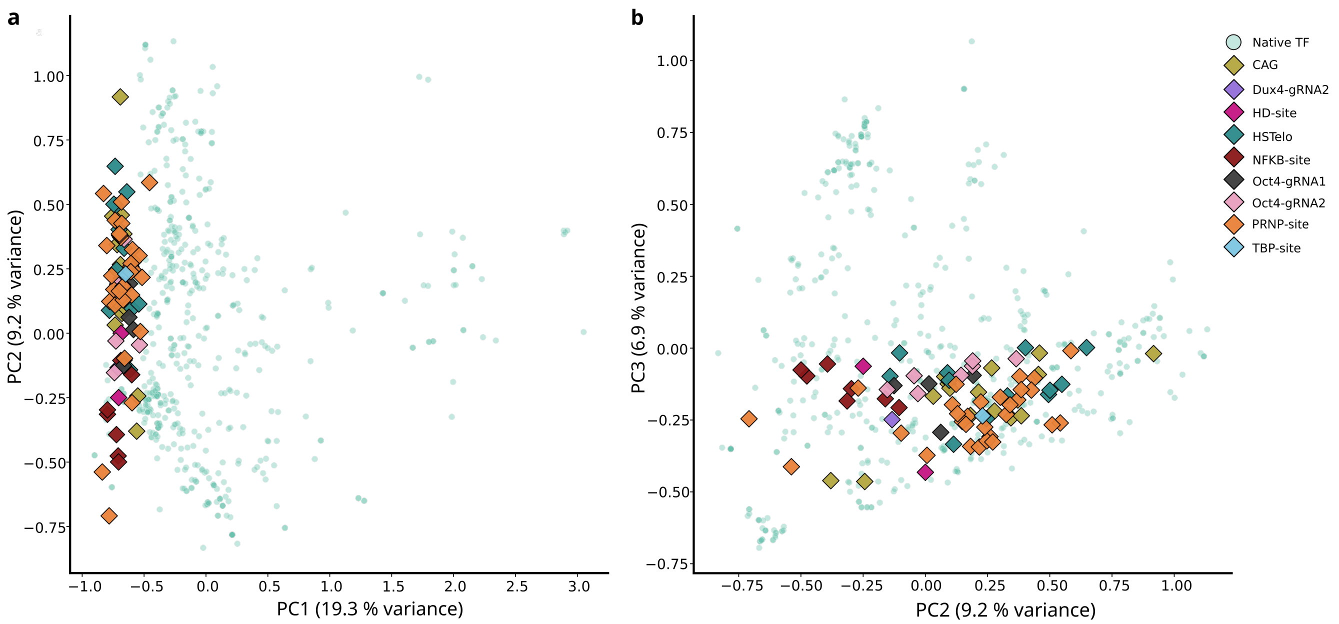
a
b
PC1 (19.3 % variance)
PC2 (9.2 % variance)
PC2 (9.2 % variance)
PC3 (6.9 % variance)
Native TF
CAG
Dux4-gRNA2
HD-site
HSTelo
NFKB-site
Oct4-gRNA1
Oct4-gRNA2
PRNP-site
TBP-site

**Fig. S10 *(previous page)*. Designed DNA binders embedded within the sequence space of native transcription factors.** PCA of ESM-2[68] sequence embeddings for protein sequences of chains extracted from crystal structures of natural protein-DNA complexes from the PDB[29,30] (teal, n = 532) and *de novo* designed DNA binding proteins (diamonds, colored by target site, n = 73). The PCA was performed on the native sequences; designed sequences were projected onto the same axes. **a,** PC1 versus PC2. **b,** PC2 versus PC3.

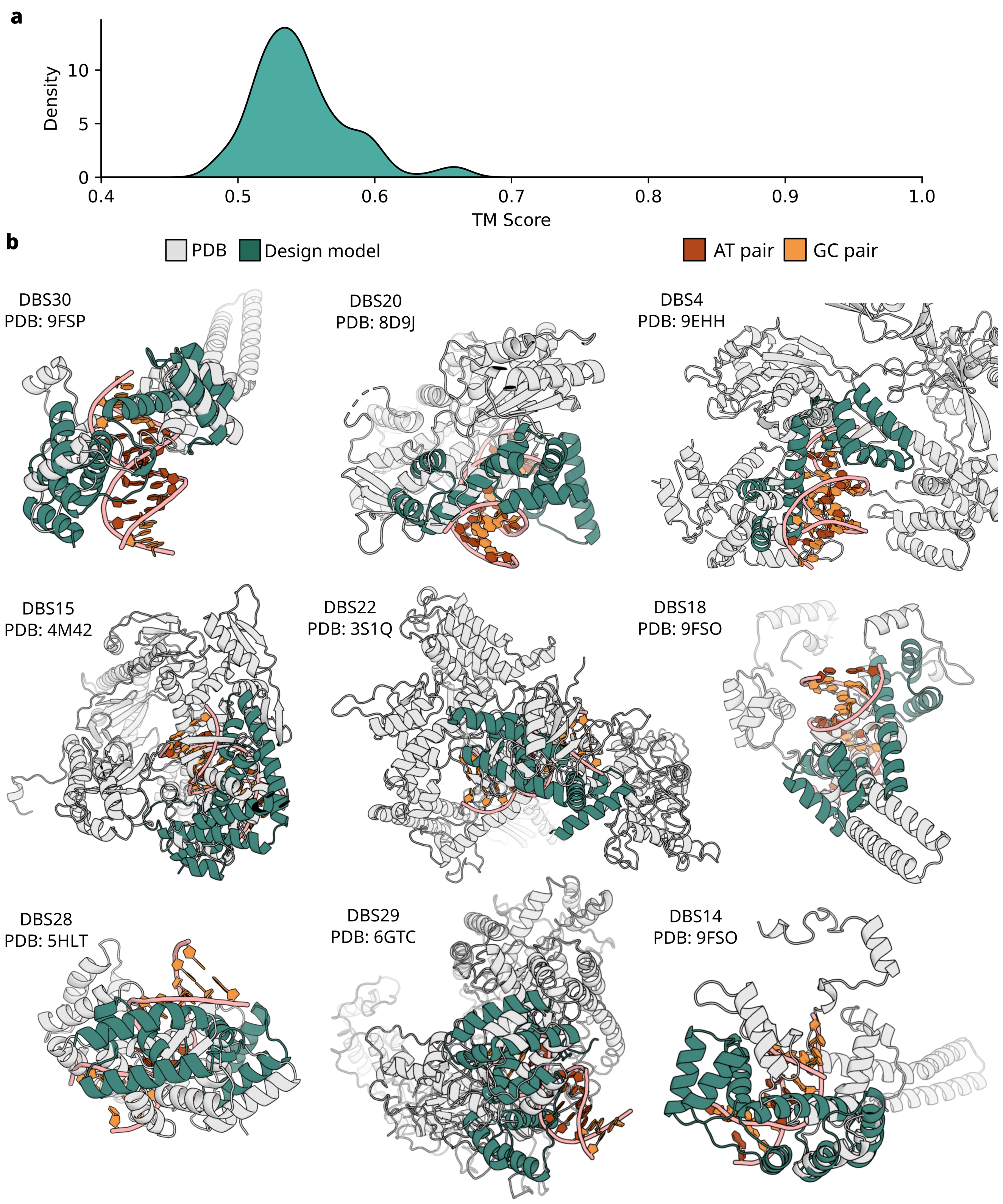
a
Density
10
5
0
0.4
0.5
0.6
0.7
0.8
0.9
1.0
TM Score
b
PDB
Design model
AT pair
GC pair
DBS30
PDB: 9FSP
DBS20
PDB: 8D9J
DBS4
PDB: 9EHH
DBS15
PDB: 4M42
DBS22
PDB: 3S1Q
DBS18
PDB: 9FSO
DBS28
PDB: 5HLT
DBS29
PDB: 6GTC
DBS14
PDB: 9FSO

**Fig. S11** ***(previous page)*****. Structural similarity of designed DNA binding proteins to native protein–DNA complexes. a,** TM-align[80] scores computed between a set of sequence-selective *de novo* DNA binders (DBs) and a reference set of protein–DNA complexes from the PDB[29,30]. For each design, the highest TM-score (best structural match) is shown as a kernel density plot. **b,** Overlays of a DB and the PDB corresponding to its best TM-align score (aligned using PyMOL[83]).

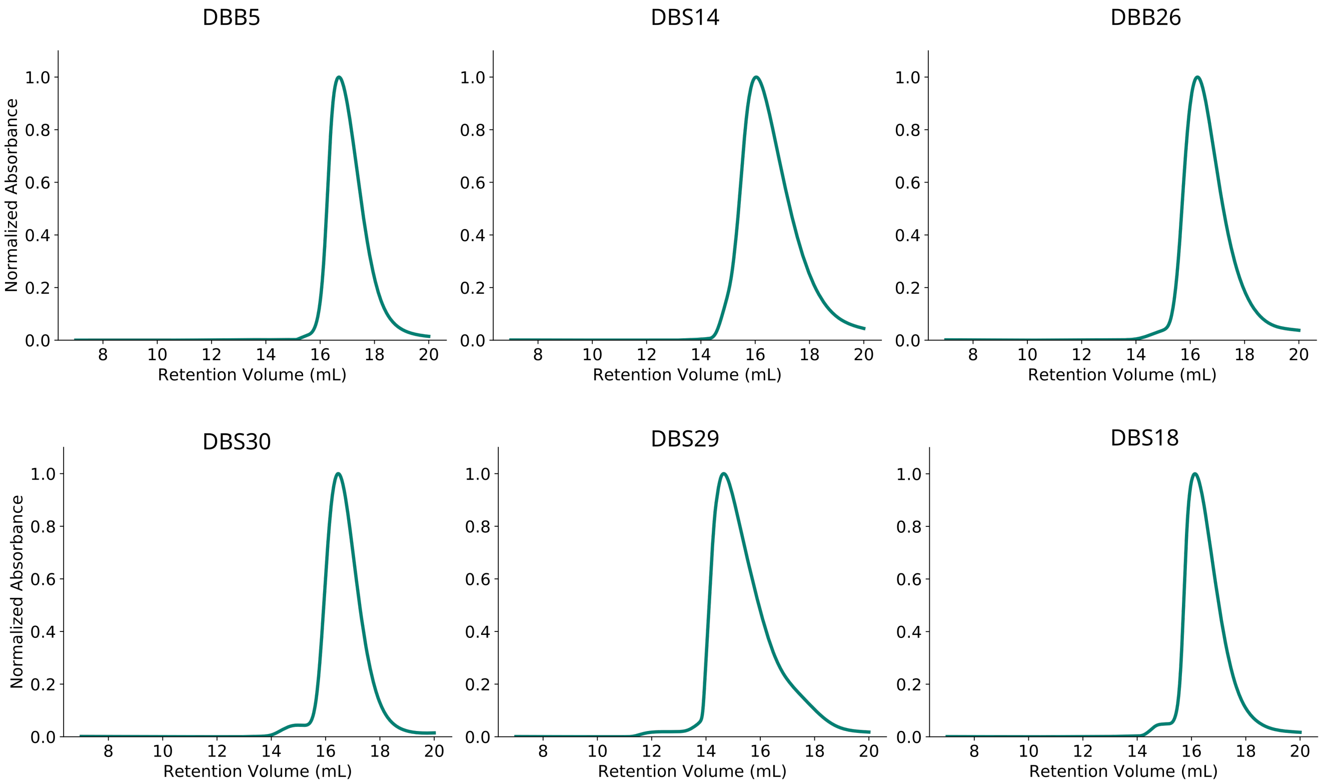
DBB5
DBS14
DBB26
DBS30
DBS29
DBS18
Normalized Absorbance
Retention Volume (mL)

**Fig. S12** ***(previous page).*** **Size-exclusion chromatography analysis of designed DNA binding proteins.** Normalized absorbance at 280nm for protein elution from a Superdex 75 Increase 10/300 GL column, indicating monodispersity of the designed DNA binders. Each trace corresponds to an individual protein sample following IMAC purification and SNAC tag cleavage[84], yielding tag-free designs.

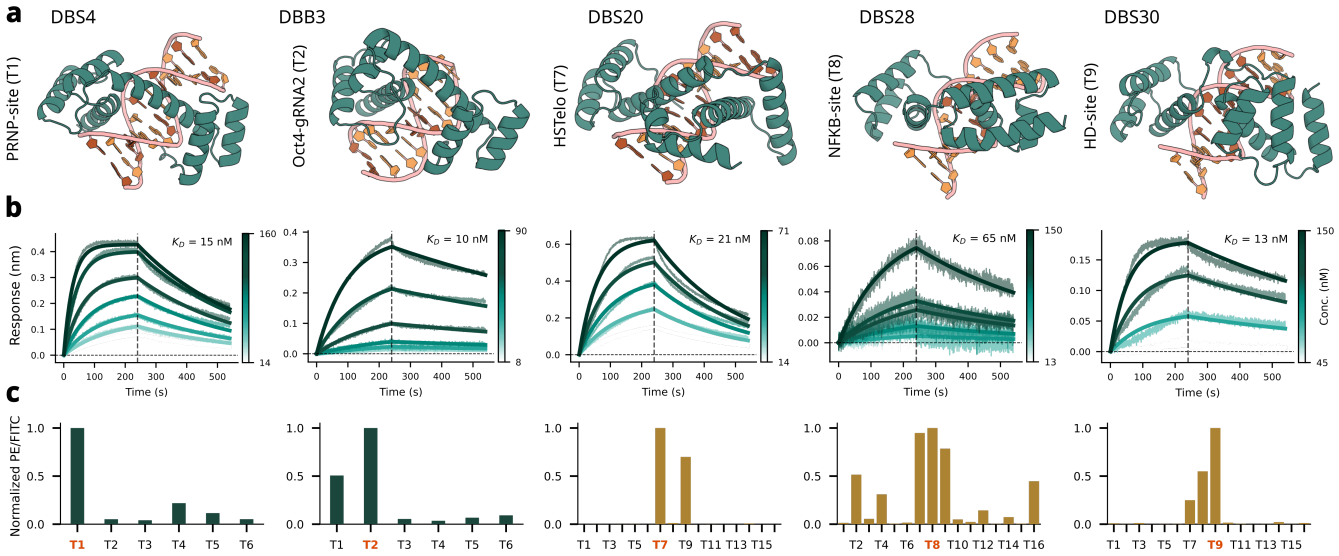

a
DBS4
DBB3
DBS20
DBS28
DBS30
PRNP-site (T1)
Oct4-gRNA2 (T2)
HSTelo (T7)
NFKB-site (T8)
HD-site (T9)
b
Response (nm)
Time (s)
Conc. (nM)
$K_D$ = 15 nM
$K_D$ = 10 nM
$K_D$ = 21 nM
$K_D$ = 65 nM
$K_D$ = 13 nM
c
Normalized PE/FITC

**Fig. S13** ***(previous page)*****. Additional structures and on-target binding affinities. a,** Structure of a representative protein-DNA complex. **b,** Bio-layer interferometry (BLI) binding kinetics for selected designs. Sensorgrams show association and dissociation across a titration series of protein concentrations. Equilibrium dissociation constants ($K_D$) were determined by global fitting to a 1:1 binding model using Octet Analysis Studio. **c,** Specificity profiles for selected designs, showing normalized binding signal (PE/FITC) across DNA targets. The intended on-target sequence for each design is indicated in orange on the y-axis (see Table 1 for target annotations). Teal bars represent hits from the core target set and yellow represents the hits for the additional target set.

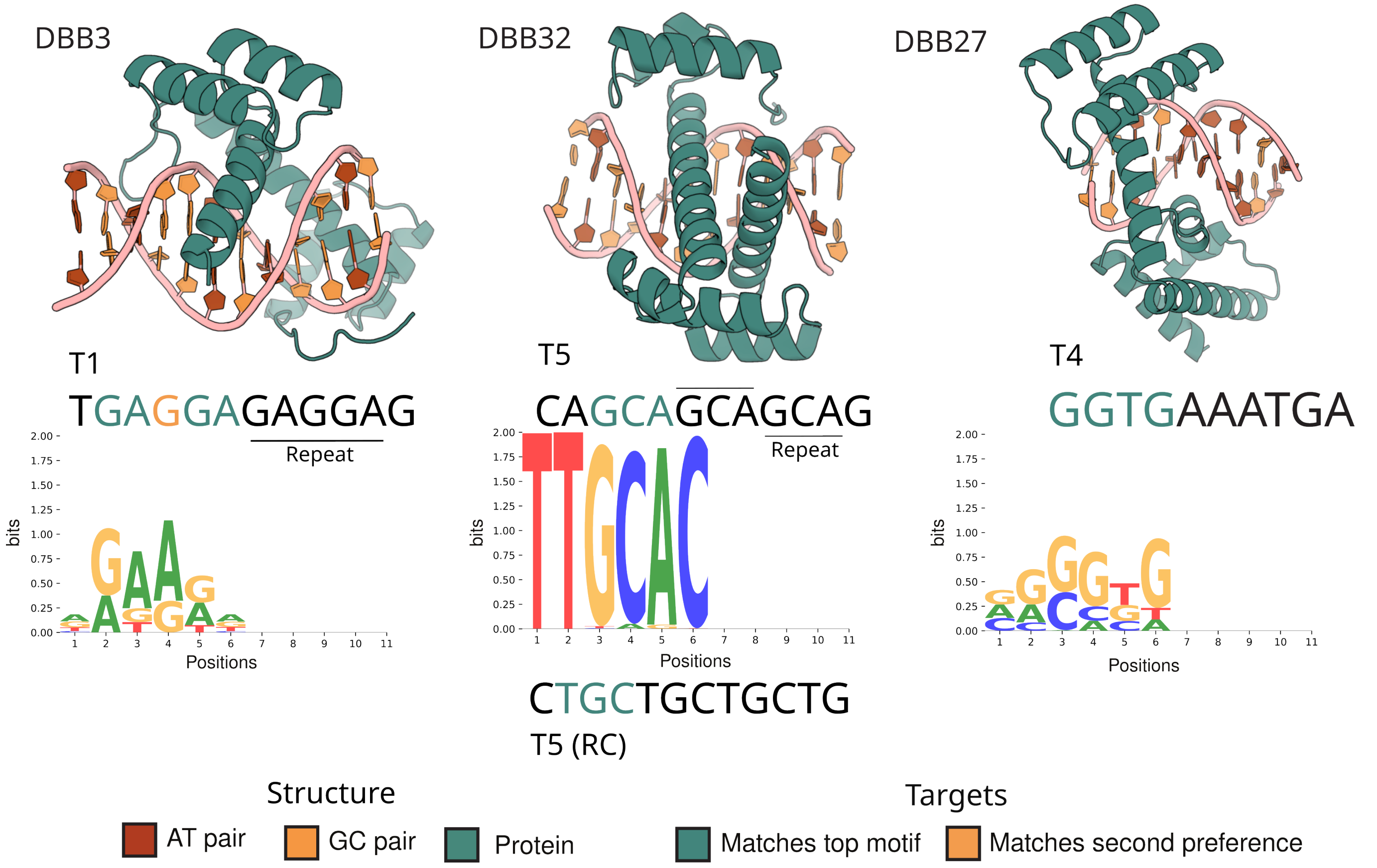
DBB3
DBB32
DBB27
T1
TGAGGAGAGGAG
Repeat
T5
CAGCAGCAGCAG
Repeat
T4
GGTGAAATGA
bits
Positions
CTGCTGCTGCTG
T5 (RC)
Structure
AT pair
GC pair
Protein
Targets
Matches top motif
Matches second preference

**Fig. S14** ***(previous page)*****. HT-SELEX characterization of binder block designs. (Top of each row)** Design models of binder block designs (DBBs) quantified via HT-SELEX[85,79] with information content-rich specificity profiles replicated across experiments. **(Bottom of each row)** PWMs derived via HT-SELEX in context of the intended target sequences for each design. Green represents a match with the intended target. Orange represents intended bases where they appear as the second preference in the HT-SELEX derived PWM. The PWMs were derived from the count matrices with an uniform background assumption.

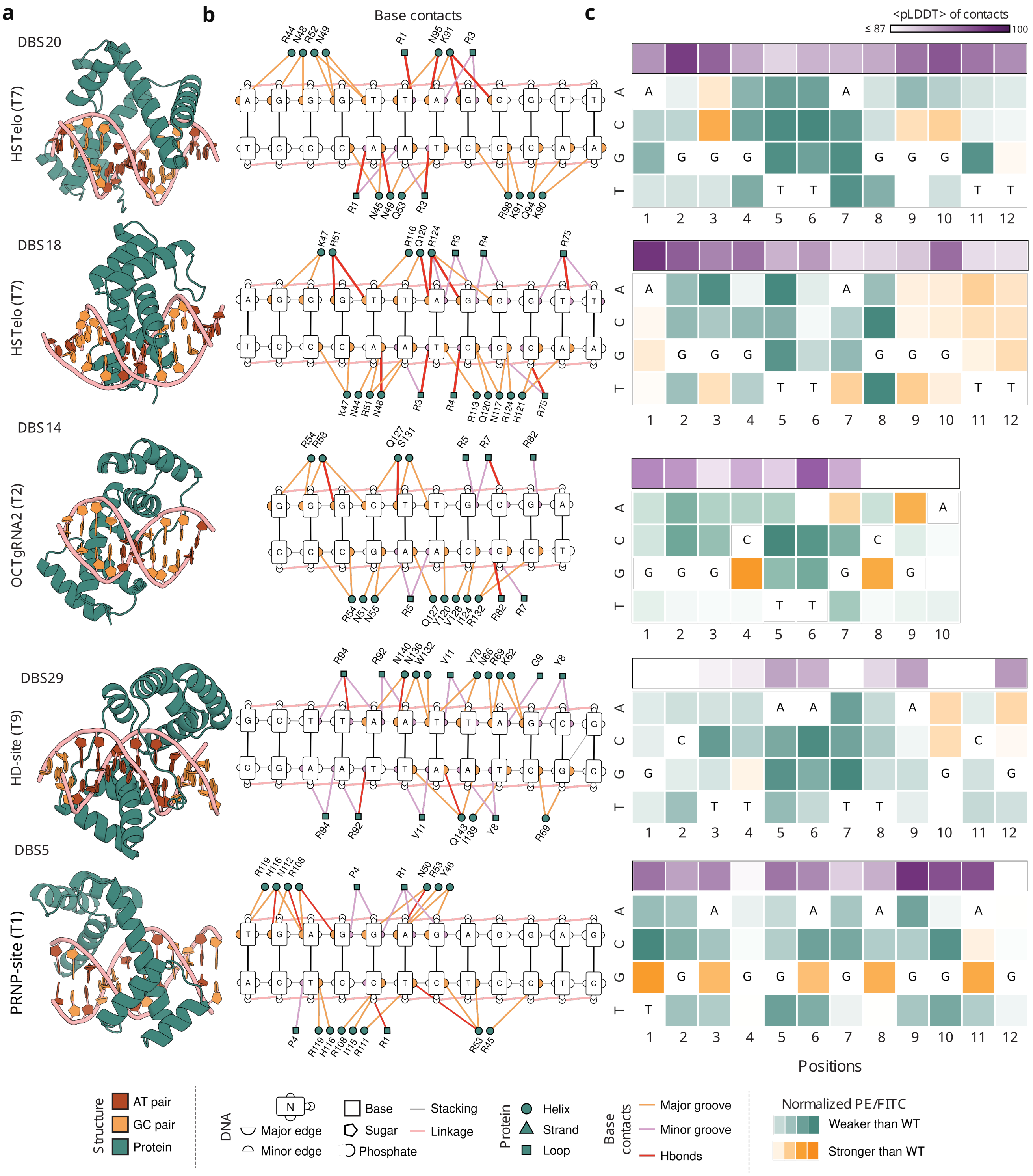
a
b
Base contacts
c
<pLDDT> of contacts
≤ 87
100
DBS20
HSTelo (T7)
DBS18
HSTelo (T7)
DBS14
OCTgRNA2 (T2)
DBS29
HD-site (T9)
DBS5
PRNP-site (T1)
Positions
Structure
AT pair
GC pair
Protein
DNA
Major edge
Minor edge
Base
Sugar
Phosphate
Stacking
Linkage
Protein
Helix
Strand
Loop
Base contacts
Major groove
Minor groove
Hbonds
Normalized PE/FITC
Weaker than WT
Stronger than WT

**Fig. S15** ***(previous page)*****. Detailed base contacts of designs evaluated by variant competition assay. a,** Design models (AF3 predictions) of selected specificity block designs characterized for fine-grained specificity using yeast surface display-based competition assay, assessing binding towards all possible single nucleotide variants of corresponding DNA target. **b,** Schematic representation of DNA base contacts (generated using DNAproDB[32] and modified stylistically) showing all protein residue contacts (with secondary structure annotation: helix, sheet, loop) to the major groove (orange) and minor groove (purple) side of the DNA bases. All Hydrogen bonds to the bases are highlighted in red. DNA backbone contacts are not shown. **c,** Results of competition assay (normalized PE/FITC) for each design (orange-green heatmap) presented in context of average AF3 pLDDT of contacting protein residues (within 4 Å) per base-pair position (purple heatmap). Green indicates a mutant competitor target was weaker than the intended (WT) base at that position, while orange indicates higher binding for the mutant competitor.

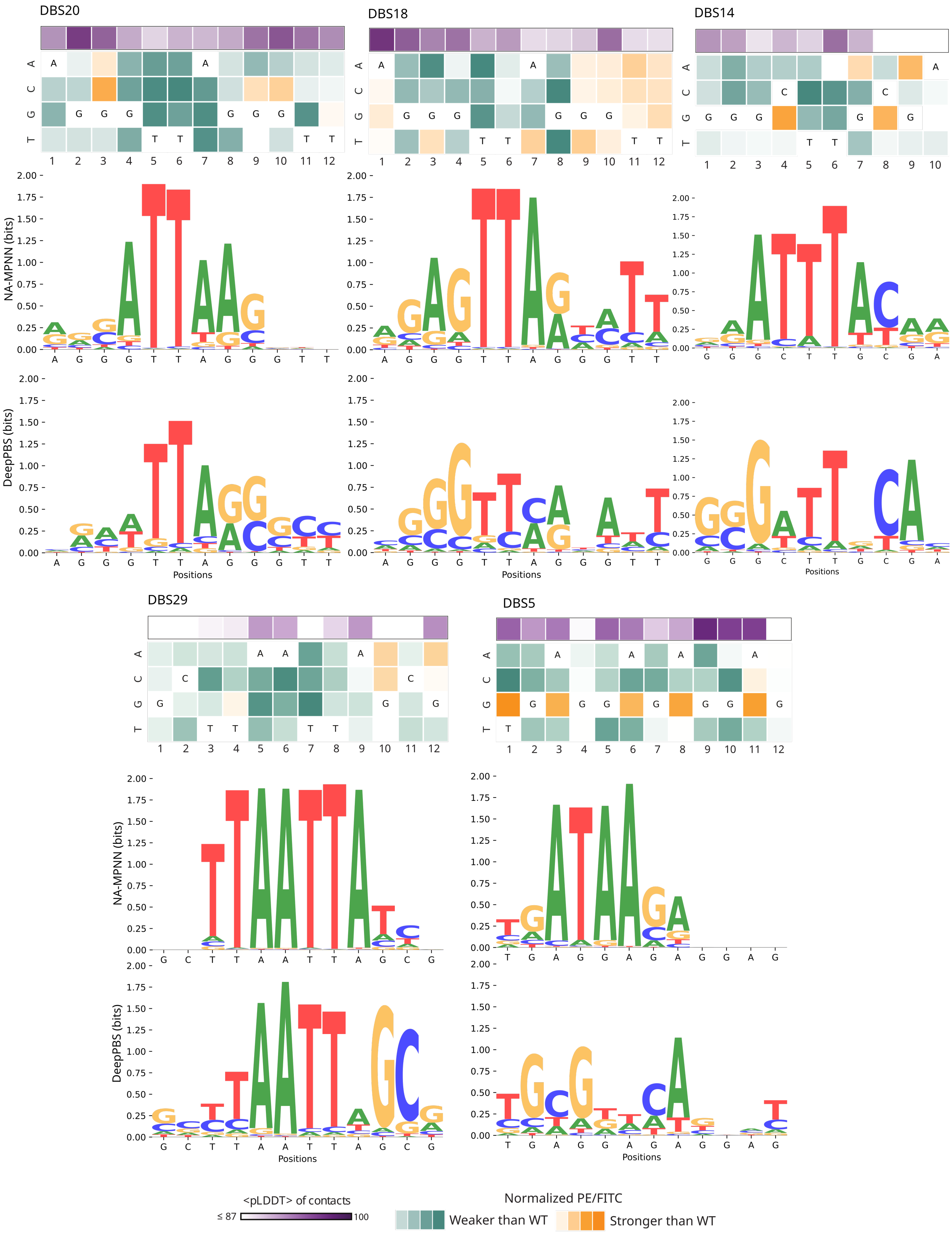
DBS20
DBS18
DBS14
DBS29
DBS5
NA-MPNN (bits)
DeepPBS (bits)
Positions
<pLDDT> of contacts
≤ 87
100
Normalized PE/FITC
Weaker than WT
Stronger than WT

**Fig. S16** ***(previous page).*** ***In-silico*** **specificity prediction.** *In silico* specificity prediction of selected designs using fixed-dock specificity prediction methods NA-MPNN[8] and DeepPBS[9]. Predicted PWMs largely corroborate the observed mutational scanning data for designs including affirming some of the off-targets effects. NA-MPNN predictions were performed with the publicly available specificity model and DeepPBS predictions were performed with the webserver. In either case, the displayed logo corresponds to information contents computed with a uniform background using the output position probability matrices.

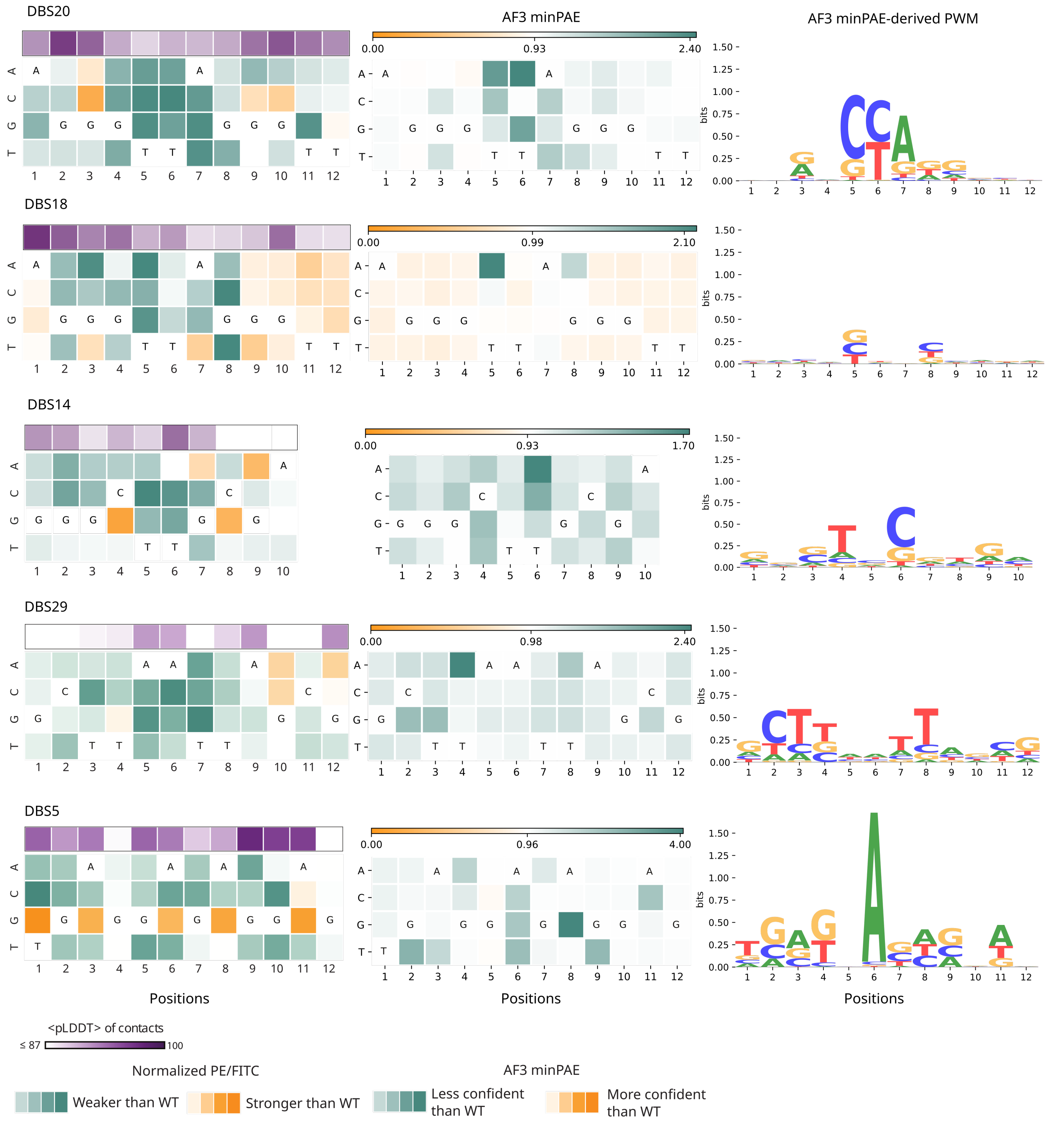
DBS20
DBS18
DBS14
DBS29
DBS5
AF3 minPAE
AF3 minPAE-derived PWM
bits
Positions
Positions
Positions
<pLDDT> of contacts
≤ 87
100
Normalized PE/FITC
Weaker than WT
Stronger than WT
AF3 minPAE
Less confident than WT
More confident than WT

**Fig. S17** ***(previous page)*****. Comparison of AF3 minPAE values with variant competition assay outcome.** Left column presents the competition assay data as presented in Fig. 4; Middle column presents AF3 output minPAE value[20] when each target variant is folded with the designed protein by AF3 (with seed 42, not templated). Wild type or intended target being white, less confident predictions in green (higher minPAE) and more confident predictions in orange (lower minPAE). The right column attempts to turn these minPAE observations into a specificity PWM. This is achieved by position-wise normalizing (via division of sum) the inverse fifth power of the minPAE values. A high power was necessary to discern any visible information content.

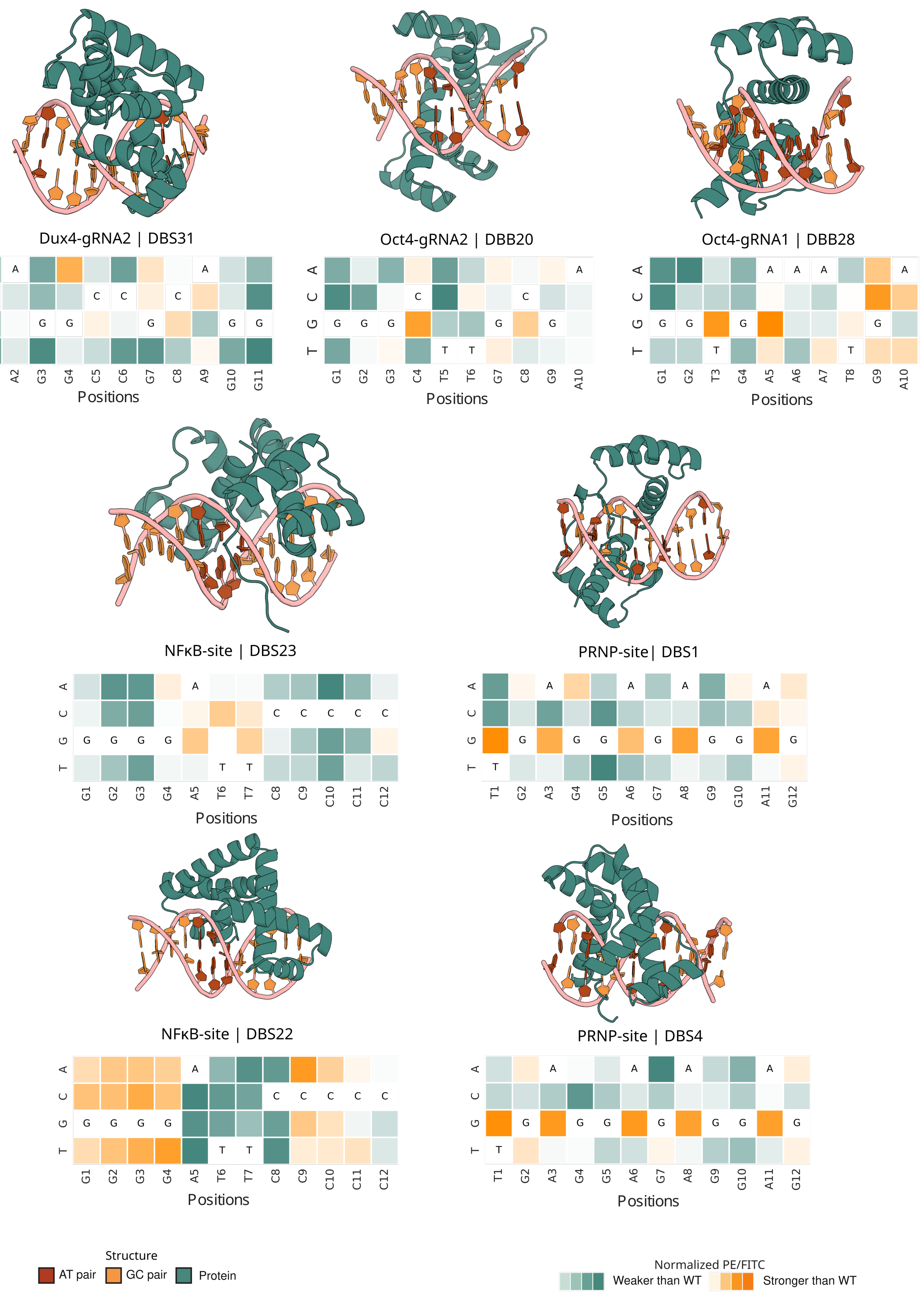
Dux4-gRNA2 | DBS31
Positions
Oct4-gRNA2 | DBB20
Positions
Oct4-gRNA1 | DBB28
Positions
NFκB-site | DBS23
Positions
PRNP-site| DBS1
Positions
NFκB-site | DBS22
Positions
PRNP-site | DBS4
Positions
Structure
AT pair
GC pair
Protein
Normalized PE/FITC
Weaker than WT
Stronger than WT

**Fig. S18 *(previous page)*. Characterization of further designs by variant competition assay. (Top of each row)** Design models (AF3 predictions) of seven further designs characterized for fine-grained specificity using yeast surface display-based competition assay, assessing binding towards all possible single nucleotide variants of corresponding DNA target. **(Bottom of each row)** Results of competition assay (normalized PE/FITC) for each design (orange-green heatmap) presented in context of average AF3 pLDDT[20] of contacting protein residues (within 4 Å) per base-pair position (purple heatmap). Green indicates a mutant competitor target was weaker than the intended (WT) base at that position, while orange indicates higher binding for the mutant competitor.

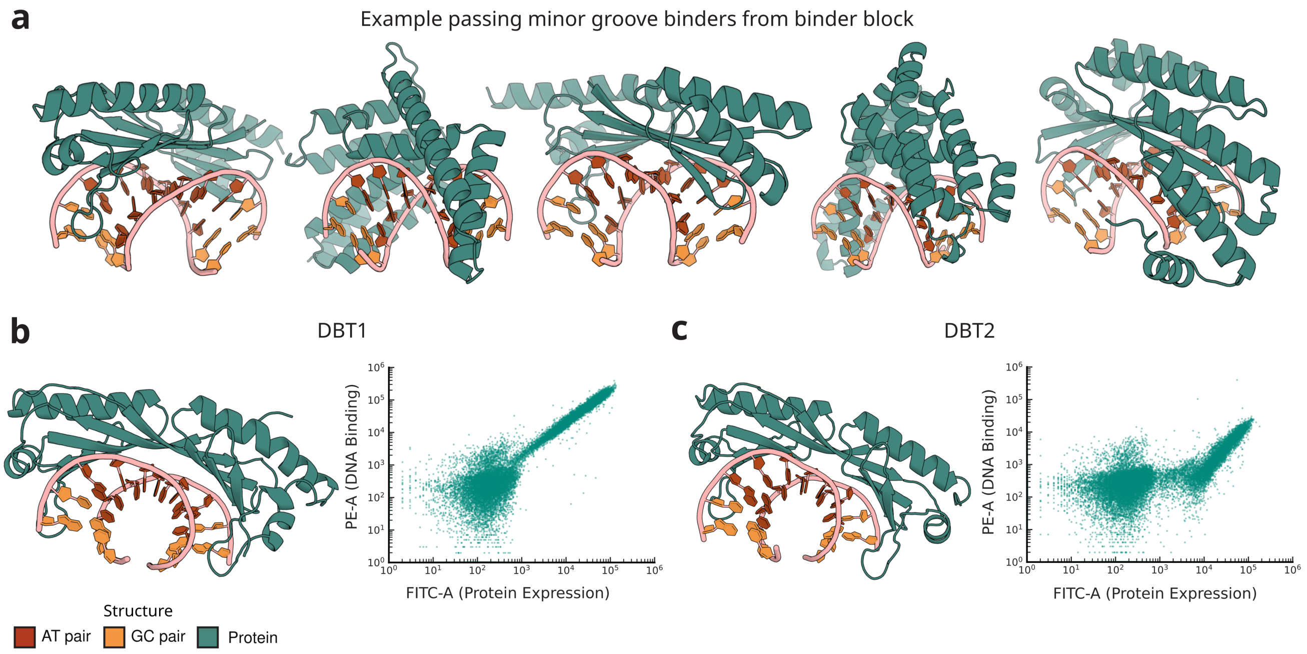
a
Example passing minor groove binders from binder block
b
DBT1
c
DBT2
PE-A (DNA Binding)
FITC-A (Protein Expression)
PE-A (DNA Binding)
FITC-A (Protein Expression)
Structure
AT pair
GC pair
Protein

**Fig. S19** ***(previous page)*****. Proof-of-principle design of minor groove DNA binding proteins. a,** Designs derived from a slightly modified design pipeline targeting a bent TBP-site target. These designs are passing designs output from the binder block of the pipeline. **b,** DBT1, a design (model at left) that was found to be a binder via yeast surface display assay (right). This design has a best case 38% sequence similarity to PDB[29,30]. **c,** DBT2, a design (model at left) that was found to be a binder via yeast surface display assay (right). This design has a best case 41% sequence similarity to PDB[29,30].

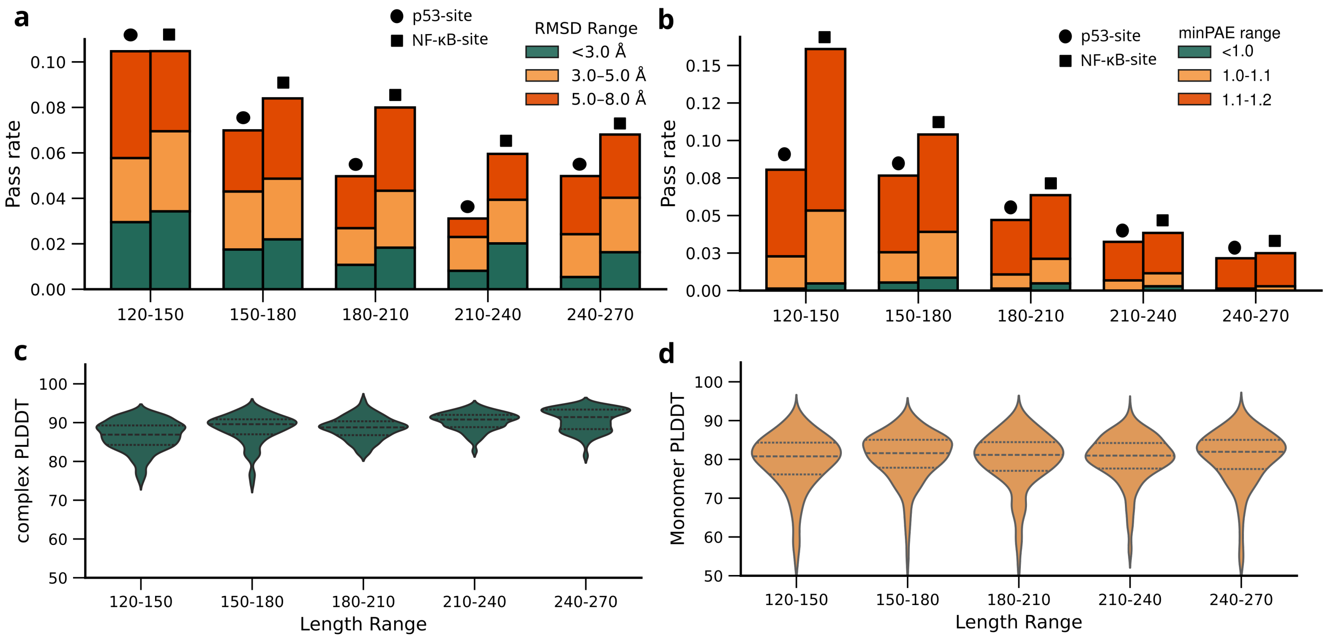
a
p53-site
NF-κB-site
RMSD Range
<3.0 Å
3.0–5.0 Å
5.0–8.0 Å
Pass rate
0.00
0.02
0.04
0.06
0.08
0.10
120-150
150-180
180-210
210-240
240-270
b
p53-site
NF-κB-site
minPAE range
<1.0
1.0-1.1
1.1-1.2
Pass rate
0.00
0.03
0.05
0.08
0.10
0.12
0.15
120-150
150-180
180-210
210-240
240-270
c
complex PLDDT
50
60
70
80
90
100
120-150
150-180
180-210
210-240
240-270
Length Range
d
Monomer PLDDT
50
60
70
80
90
100
120-150
150-180
180-210
210-240
240-270
Length Range

**Fig. S20 *(previous page)*. Sampling longer lengths of protein.** For two design targets (p53-site and NF$\kappa$B-site), a small scale design cycle was carried out (RFD3 diffusion[18], LigandMPNN[19], sequence design AF3[20] folding) for varying length ranges. All hyperparams were kept as described in (see Methods) except for a reduced `n_batches` 15 and `diffusion_batch_size` of 2. **a,** Pass rates of designs based on different thresholds of DNA-aligned protein RMSD. Longer lengths result in lower pass rates but still at a viable 5-6% at length 240-270. **b,** Pass rates based on different AF3 minPAE[20] threshold showing drop over larger lengths, although at the most stringent cut-off ($< 1.0$) length range 150-180 performs the best. **c,** Average AF3 pLDDT[20] of protein in complex with DNA, (for designs with $< 5$ Å DNA-aligned protein RMSD) over different length ranges show an increasing trend. **d,** AF3 predictions of protein only (without DNA) for the same set used in (**c**) show no change in pLDDT distribution over different length ranges.

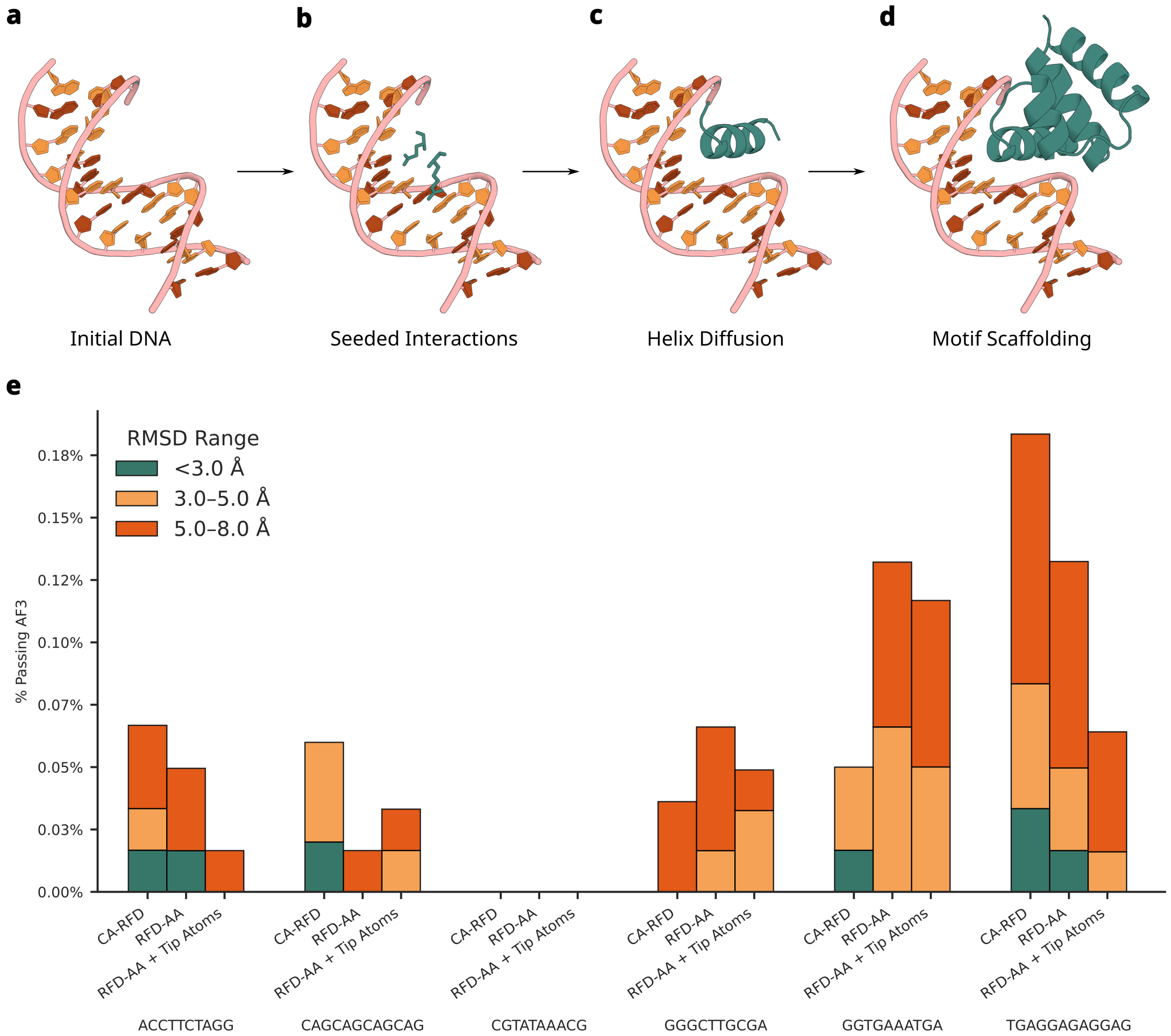
a
b
c
d
Initial DNA
Seeded Interactions
Helix Diffusion
Motif Scaffolding
e
RMSD Range
<3.0 Å
3.0–5.0 Å
5.0–8.0 Å
% Passing AF3
0.18%
0.15%
0.12%
0.10%
0.07%
0.05%
0.03%
0.00%
CA-RFD
RFD-AA
RFD-AA + Tip Atoms
ACCTTCTAGG
CAGCAGCAGCAG
CGTATAAACG
GGGCTTGCGA
GGTGAAATGA
TGAGGAGAGGAG

**Fig. S21** ***(previous page)*****. RFdiffusion All-Atom pipeline and *in silico* results.** To generate backbones with RFdiffusion All-Atom (RFD-AA), we apply the following pipeline. **a,** First, we start from a predicted DNA structure. **b,** We then use RIFGen[86] to sample amino acid residues aligned to DNA base atoms in the major groove of the DNA. RIFGen samples these residues from the set of amino acids found in native DNA binding proteins in the PDB[29,30] that are making hydrogen bond contacts to DNA bases, and are therefore likely to feature biophysically plausible interaction geometries. **c,** Tip-atoms (hydrogen bond contact atoms) from these amino acid residues, along with the DNA structure are used as inputs to a finetuned RFdiffusion All-Atom[24] see supplementary information, to generate a small helix in the major groove of the DNA. The tip-atoms serve to condition the model toward generating a helix with amino acid residues that can make good contacts to the DNA. **d,** Helices that have good *in silico* properties are then motif scaffolded to build the rest of the protein backbone. **e,** Backbones were generated using the CA-RFD model, RFD-AA model, and with the full tipatom RFD-AA pipeline, against the six core DNA targets described in Fig. 1. The designs are for 50-80 amino acid length proteins, which was an earlier design goal. After designing 5 sequences per backbone with LigandMPNN, the fraction of sequences passing AF3 RMSD thresholds for each model and DNA target sequence is reported.

### Supplementary Information

#### minPAE as filter for specificity for native transcription factors

The minimum interchain PAE value in AF3 predictions (minPAE) has shown good performance for *in silico* filtering of binders designed against protein targets[87]. Let, P be the set of protein residues and D be the set of DNA residues. We defined minPAE for protein-DNA pairs as,

$$minPAE = min_{i \in P, j \in D} PAE(i, j)$$

We evaluated minPAE's ability to discriminate on-target from off-target sequences by running AF3 predictions of native DNA binding proteins (DBs) in complex with their target sequences and with randomly mutated sequences. We performed AF3 predictions of a subset of transcription factors (TFs) contained in the Uniprobe database[81] interacting with their recorded consensus sequences and mutated sequences derived from them. minPAE presented a moderate signal to discriminate binder sequences (PWM score $> 0$) from non-binders with a mean per-TF AUC of 0.69, independently of the use of MSA information (Fig. S3a,b). Good performance in single-sequence mode prediction is essential for *de novo* designed binder filtering due to their lack of evolutionary sequence information. We also benchmarked minPAE on a mutational library of the Saccharomyces cerevisiae bHLH transcription factor Pho4 binding to variants of its cognate E-box, where affinity is quantified as an apparent $K_D$[82]. Defining binders as DB–DNA pairs with $K_D < 10\ \mu M$, minPAE discriminated binders from non-binders with a mean per-sequence AUC of 0.75 when AF3 was run without an MSA, but collapsed to random performance, 0.51, when MSA information was provided (Fig. S3c,d). We interpreted that addition of the MSA evolutionary constraints reduces the contribution of subtle geometric and chemical interface differences between point mutants, preventing minPAE from discriminating binding from non binding DNA-Pho4 pairs. Pairs correctly flagged as likely non-binders by high minPAE values shift to low minPAE when adding the MSA information, producing false positives (Fig. S3e). Together, these two benchmarks show that minPAE is a useful *in silico* predictor of DB–DNA binding, and that its discriminative power is increased in the single-sequence regime.

#### Diversity of designs

To quantify the uniqueness of our designs, we computationally embedded our design targets and native TF target sites from the JASPAR database[60] (see main:MethodsMethods) in an embedding space computed via a DNA language model, Evo[67] (see Methods). Our targets span a broad range of DNA sequence space and some of the successful targets appear to be distinct from the core TF target sequence space (Fig. 2g). This demonstrates the potential of this approach for targeting unique sites. We also performed a similar analysis for the designed protein sequences. We project the designed sequences into an ESM-2-based[68] embedding space of TF sequences derived from crystal structures (see Methods). We find that the designed proteins occupy a distinct cluster separated from the natives and demonstrate a broad range within the cluster (Fig. S10). To quantify structural similarity of our designs to the Protein Data Bank (PDB)[29,30], we performed TM-align[80] of each design against structures in the PDB which returned best matches of TM-scores around 0.5-0.6 (Fig. S11a). Overlays with closest matches are displayed in Fig. S11b.

#### HT-SELEX characterization of binder block designs

Specificity of native TFs are often quantified by screening against a randomised library of DNA targets, establishing the broadest picture of specificity[88,79,89,90,91]. TF binding data from such experiments show considerable binding towards multiple sites[92] ranging from fairly promiscuous (e.g. HoxA5 JASPAR entry MA0158.2) to very specific (e.g. Max dimer, JASPAR entry MA0058.4). We carried out HT-SELEX experiments (see Methods) on several binder block designs (DBBs) (Fig. S14). Design DBB5 shows strong specificity towards an 8-mer PWM with 5 positions aligning with the intended target PRNP-site (Fig. S14). DBB3, another binder towards PRNP-site target shows a 5-mer PWM with close correspondence to positions 2-6 (GAGGA) of intended target (this 5-mer repeats again in this target at positions 7 to 11) (Fig. S14). DBB32, a binder towards the CAG-repeat target strongly recognizes the repeating unit GCA, overlapping with the reverse complement repeating unit TGC, which is a GCA offsetted by one base in the opposite strand (Fig. S14) consistent with potential dimeric or multimeric binding. DBB31, also targeting CAG-repeat, has a longer binding profile with more modest information content, matching the intended target as for either top or second-best preference for seven positions. DBB26 targeting the AT-rich target TBP-site shows a 10-mer PWM, with 7 out of 10 positions of the top motif matching the intended target (Fig. S14). DBB20, design targeting the OCT4-gRNA2 also has a 10-mer motif with 6 out 10 positions closely resembling the intended target. Lastly, DBB27, targeting OCT4-gRNA1 shows a shorter binding motif of length 6 with 4 positions closely matching the intended target (Fig. S14). In summary, although not screened *in silico* for broad specificity, we observe information content-rich binding signals for multiple binders, with patterns matching the intended target to varying degrees (Fig. S14).

#### Minor groove binder design targeting non-standard DNA conformation

To design minor groove binding proteins presented in Fig. S19 we used a modified version of the binder block (Fig. S1). Target structure was obtained by AF3[20] folding the TBP-site target (CGTATAAACG) with the native TF. Modifications for input to RFD3[18]: diffused protein lengths 150-220, ORI placement perpendicularly to the local helical axis and 1.5 Å into the minor groove direction, one ORI per 8-mer, hydrogen bond annotations on respective minor groove base atoms. First round LigandMPNN[19] sampled 10 sequences per backbone. AF3 pass rates were much lower for this case: 0.67%. Second round LigandMPNN sampled 50 sequences. Final filtering cutoffs: DNA-aligned protein RMSD 3Å, ipTM 0.7, total hydrogen bond: 14, hydrogen bond support count (including protein backbone hydrogen bonds) : 10, minPAE: 1.25, minor groove hydrogen bond count 2. A total of 46 designs passed these criteria and were validated experimentally leading to 2 successful binders.

#### CA-RFdiffusion finetuning

We adapt the C-alpha diffusion and refinement framework in Lauko et. al.[23] for the task of DNA binder design. For the diffusion model, we finetune the protein-nucleic acid complex prediction model RoseTTAFoldNA[93] for the diffusion generative task, on a subset of the PDB[29,30] containing only protein-DNA interfaces cropped tightly around a protein contact with either a DNA base or backbone atom. By not backpropagating a loss on the frame rotations of the backbone frames, this converts RFdiffusion to a C-alpha only model, generating only C-alpha coordinates. A subsequent refinement model is trained to recover the correct backbone frames in a single step after the alpha carbon backbone framework has been generated by the diffusion model.

#### RFdiffusion All-Atom finetuning

We adapt the tip-atom motif scaffolding training framework in Krishna et. al.[24] to generate DNA binders. The model was finetuned starting from RosettaFold All-Atom structure prediction weights on the diffusion generative task. The training data included both native protein-nucleic acid complexes in the PDB[29,30] and a distillation set of predicted transcription factor structures from Baek et. al.[93]. The structures were cropped around a protein contact with either a DNA base or backbone atom, to train the model to learn protein-DNA interface structures. The model was trained on both the tip-atom placement task, where DNA contacting protein side-chains are converted to their atomic representation, and the model is given the location of the contacting tip-atoms but must recapitulate the rest of the side-chain, and the motif scaffolding task, where the model is given the backbone coordinates of a contiguous stretch of protein close to the DNA and must recapitulate the remaining backbone scaffold. Radius-of-gyration, percent helicity, and center-of-mass conditioning was also applied to give more fine-grained control of desired design parameters.

[30] wwPDB consortium. Protein Data Bank: the single global archive for 3D macromolecular structure data. *Nucleic acids research*, 47(D1):D520–D528, 2019.

[31] Luca Mariani, Xiao Liu, Kwangwoon Lee, Stephen S Gisselbrecht, Philip A Cole, and Martha L Bulyk. Dna bendability regulates transcription factor binding to nucleosomes. *Nature Structural & Molecular Biology*, 32(11):2185–2195, 2025.

[32] Raktim Mitra, Ari S Cohen, Jared M Sagendorf, Helen M Berman, and Remo Rohs. Dnaprodb: an updated database for the automated and interactive analysis of protein–dna complexes. *Nucleic acids research*, 53(D1):D396–D402, 2025.

[58] Eric Zhao, Pranjal Awasthi, and Sreenivas Gollapudi. Sample, scrutinize and scale: Effective inference-time search by scaling verification. *arXiv preprint arXiv:2502.01839*, 2025.

[59] Xiang-Jun Lu, Harmen J Bussemaker, and Wilma K Olson. Dssr: an integrated software tool for dissecting the spatial structure of rna. *Nucleic acids research*, 43(21):e142–e142, 2015.

[60] Ieva Rauluseviciute, Rafael Riudavets-Puig, Romain Blanc-Mathieu, Jaime A Castro-Mondragon, Katalin Ferenc, Vipin Kumar, Roza Berhanu Lemma, Jérémy Lucas, Jeanne Chèneby, Damir Baranasic, et al. Jaspar 2024: 20th anniversary of the open-access database of transcription factor binding profiles. *Nucleic acids research*, 52(D1):D174–D182, 2024.

[61] Edwin N Neumann, Tessa M Bertozzi, Elaine Wu, Fiona Serack, John W Harvey, Pamela P Brauer, Catherine P Pirtle, Alissa Coffey, Michael Howard, Nikita Kamath, et al. Brainwide silencing of prion protein by aav-mediated delivery of an engineered compact epigenetic editor. *Science*, 384(6703):ado7082, 2024.

[62] Susan E Andrew, Y Paul Goldberg, Berry Kremer, Håkan Telenius, Jane Theilmann, Shelin Adam, Elizabeth Starr, Ferdinando Squitieri, Biaoyang Lin, Michael A Kalchman, et al. The

relationship between trinucleotide (cag) repeat length and clinical features of huntington's disease. *Nature genetics*, 4(4):398–403, 1993.

[81] Maxwell A Hume, Luis A Barrera, Stephen S Gisselbrecht, and Martha L Bulyk. Uniprobe, update 2015: new tools and content for the online database of protein-binding microarray data on protein–dna interactions. *Nucleic acids research*, 43(D1):D117–D122, 2015.

[82] Arjun K Aditham, Craig J Markin, Daniel A Mokhtari, Nicole DelRosso, and Polly M Fordyce. High-throughput affinity measurements of transcription factor and dna mutations reveal affinity and specificity determinants. *Cell Systems*, 12(2):112–127, 2021.

[83] Schrödinger, LLC. The PyMOL molecular graphics system, version 1.8. November 2015.
